# Physiologically based demographic model/GIS analyses of thirteen invasive species in Africa: why the biology matters

**DOI:** 10.1101/2024.09.17.613432

**Authors:** Andrew Paul Gutierrez, Luigi Ponti, Markus Neteler, Jose Ricardo Cure, Peter E. Kenmore, George Simmons

## Abstract

Globally, research and policy groups often lack the expertise to develop appropriate models to analyze agroecological and invasive species problems holistically to inform management and quarantine policy development under extant and climate change over wide geographic landscapes. Off-the-shelf species distribution models (SDM) correlate weather and other variables to records of species presence and have become the mainstay for predicting the geographic distribution and favorability of invasive species (Elith 2017). However, SDM analyses lack the capacity to examine the underpinning dynamics of agroecosystems required to inform policy and develop management strategies. We propose that age-structured physiologically based demographic models (PBDMs) can solve important aspects of this challenge as they can be used to examine prospectively species dynamics locally as well as their potential geographic distribution and relative abundance across vast areas independent of presence records. PBDMs fall under the ambit of time-varying life tables (TVLTs; cf. Gilbert *et al*. 1976) and capture the weather driven biology, dynamics, and interactions of species, and can be used to examine the system from the perspective of any of the interacting species. Here, we use the PBDM structure to examine the dynamics across Africa of thirteen invasive species from various taxa having diverse biology and trophic interactions (see Gutierrez 1996, Gutierrez and Ponti 2013a). PBDMs are perceived to be difficult to develop, hence the *raison d’être* is to show this is not the case and illustrate their utility invasive and endemic agricultural and medical/veterinary pest species at the local and the large geographic scale of Africa. We note that PBDMs provide a structure for continued model improvements.

The development of open access software is proposed to facilitate PBDM development by non-experts emphasizing the crucial role of sound biological data on species responses to weather and to other species in a multi-trophic, interactions, and provide a guide for collecting the appropriate biological data. While the emphasis is on plant/arthropod interactions, models of diseases can be accommodated. The Supplemental Materials summarizes a large array of heritage PBDMs reported in the literature based on the methods outline herein, noting that the same model structure can be used to analyze and manage non pest species.

## Introduction

Cuthbert et al. (2022) reported that since 1960, … management expenditures have totaled at least US$95.3 billion (in 2017 values), … 12-times less than damage costs from invasions ($1130.6 billion), … the effectiveness of management expenditure is difficult to assess due to a lack of standardized measurement across spatial, taxonomic and temporal scales. Other estimates of worldwide estimated annual losses due to invasive species (FAO, 2023) include Pimentel *et al*. (2000) who opined that collectively invasive species cause more than US$140 billion in losses annually in the United States, losses of US$28 billion with projected losses of US$148 billion in the European Union by 2040 (Henry *et al*. 2023). Estimated yield loss to subsistence farmers from pest/disease species in Africa are large but not well documented (Pyšek *et al*. 2008, Khuroo *et al*. 2011) where pests/diseases reduce yields of basic food staples (e.g., maize, millet, sorghum), and vectored tropical diseases affect the health of millions of people. The focus of this study is Africa.

The estimates of losses are large and beg serious attention, but how to analyze agroecological and invasive species problems holistically to inform management and quarantine policy development under extant and climate change over wide geographic landscapes remains an open important question. The capacity to predict prospectively the dynamics and geographic distribution and relative abundance of pest species and their impact under current and future climatic conditions in both developed and developing economies is critical for management and policy development. Commonly, researchers study the biology of species without a unifying framework to analyze the problems resulting in myriad disjointed studies. Readily available species distribution models (SDM) that correlate weather and other variables to records of species presence are the mainstay for prospectively predicting the geographic distribution and favorability for invasive species (Elith 2017). However, SDMs lack the capacity to examine the weather driven biology and dynamics and interactions of species required to inform holistic management and policy development. We propose that weather driven physiologically based demographic models (PBDMs) circumvent many of these limitations and strongly augment SDM and other research approaches. The capacity to develop PBDMs would enable researchers and policy makers globally to assess the time varying dynamics of target species in agricultural and in natural systems independent of time and place constraints; to assist the development of strategic and tactical responses to and management of plant-herbivore-natural enemy interactions and vectored medical and veterinary disease interactions under extant and climate change weather Gutierrez 1996, Gutierrez and Ponti 2013a).

The focus of our study is to show by example how the same model structure can be used to examine the biology and dynamics of diverse species. Specifically, to make PBDM development and implementation universally accessible, we propose the development of a web application programming interface (Web API) written in a modern programming language like Python to enable users to develop and run models remotely using internet links (see Supplemental Materials appendix **Figure 1**) as is currently the case with off-the-shelf SDMs. The PBDM software would guide assembling the relevant weather driven biology of species and of species interactions, linkages to weather files to run the model, software to summarize the multi-year runs in georeferenced lattice cells across large regions (e.g., globally), and provide seamless linkages to GIS and statistical software to map and analyze the results (see **Fig. 1**). Sub-**Fig, 1E** is an example of the complexity that has been modelled using PBDMs (Gutierrez and Baumgärtner 1984, Gutierrez and Ponti 2013b). The Web API would also enable linkages to the executables of heritage models (e.g., Supplemental Materials Table 1) written in various computer languages (e.g., Fortran, Pascal, C++, Python). A Web API prototype was developed and implemented by the EU funded MED-GOLD project (https://doi.org/10.3030/776467) as a component of the project’s Information and communication technology (ICT) platform, and the heritage Pascal based olive/olive fly model (Gutierrez *et al*. 2009) was implemented and used to analyze olive production in Andalusia, Spain, and to map the results using the open-source GRASS-based GIS interface (Ponti *et al*. 2023).

**Figure 1.**
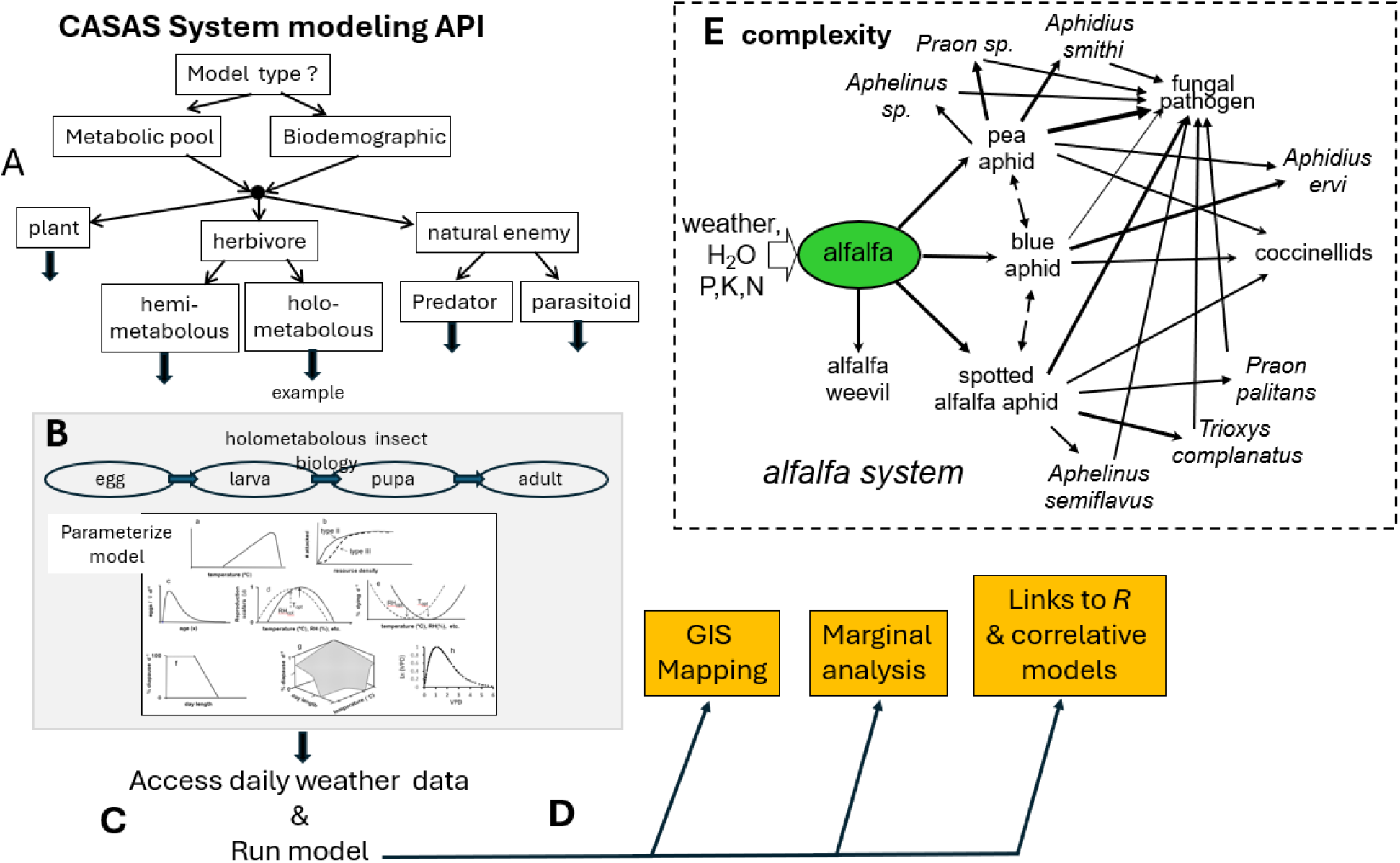
API Platform for developing and implementing physiologically based demographic models (PBDMs): **(A)** decision path for system model development; **(B)** sub model parameterization (see **Figs. 2**,**3**,**4**); **(C)** access weather files and run models; **(D)** create output files for mapping and analysis; and an example of the complexity modeled with PBDMs (e.g., Gutierrez and Baumgärtner 1984; Gutierrez and Ponti 2013c). Note in sub-**Fig. 1A**, hemi- and holometabolous life cycles may occur in any arthropod trophic level, and disease models can also be included, and parasitized aphids are also consumed by coccinellids (**Fig. 1E**).

Most of the examples reported here and in Supplemental Materials **Table 1** were developed by colleagues in the casasglobal.org consortium, and hence self-citation is unavoidable. Complete biological details for each model are found in the cited references. Last, not all the heritage models reported herein have the same level of biological detail, but all have the same underlying structure outlined below.

The focus of this paper is to illustrate how the same model structure is used to develop PBDMs for poikilotherm species dynamics and their interactions with oother species, and to use the models prospectively to predict the geographic distribution and relative abundance of system components (i.e., our metrics of favorability). If sufficiently complete, the models can be used as the objective function in economic analyses of agroecological problems (Regev *et al*. 1998, Pemsl *et al*. 2007, Gutierrez et al. 2020). The dynamics of thirteen invasive species listed below in no historical or geographic order were modeled, and their prospective ranges across Africa mapped with sub maps of other regions given as appropriate. Some of the species are agricultural pests and others are medical and veterinary pests. The biological details used to develop the PBDMs are found in the references cited for each. Note that species with a wide native geographic distribution may have region specific strains/biotypes, strains which may invade a novel environment with their behavioral and physiological traits determining establishment success, dynamics and adaptation to the new environment (see Gutierrez *et al*. in press). The PBDMs were developed using data available in the literature, and a brief statement of the limitations of some of the models. N.B. PBDMs can be easily improved as new data becomes available without changing the basic structure of the model. The species examined are the following.

(1) new world screwworm fly (*Cochliomyia hominivorax* (Coquerel))
(2) olive/olive fly (*Olea europaea* L*/ Bactrocera oleae* (Rossi))
(3) Asian brown marmorated stinkbug (*Halyomorpha halys* (Stål)) and three of its parasitoids
(4) South American tomato pinworm *Tuta absoluta* Meyrick
(5) African false codling moth (*Thaumatotibia leucotreta* (Meyrick))
(6) Fall armyworm (FAM, *Spodoptera frugiperda* (J. E. Smith))
(7) citrus/Asian citrus psyllid (*Diaphorina citri* Kuwayama)/parasitoid *Tamarixia radiata* (Waterston)/the bacterial pathogen (‘*Candidatus* Liberibacter asiaticus’)
(8) mosquito (*Aedes albopictus* (Skuse))
(9) mosquito (*Aedes aegypti* (L))
(10) Mediterranean fruit fly (*Ceratitis capitata* (Wiedemann)
(11) melon fly (*Bactrocera cucurbitae* (Coquillett))
(12) oriental fruit fly (*Bactrocera dorsalis* (Hendel))
(13) Mexican fruit fly (*Anastrepha ludens* (Loew)

### Model overview

PBDMs fall under the ambit of time-varying life tables (TVLTs; cf. Gilbert *et al*. 1976), the theoretical bases of which were reviewed by Gutierrez (1992, 1996), Gutierrez *et al*. (1994), and Mills and Gutierrez (1999). The initial impetus for the development of (PBDMs/TVLTs) was the early work of R.D. Hughes in Australia (Hughes and Gilbert 1968), N.E Gilbert in Canada (Gilbert and Gutierrez 1973) and Gutierrez and Baumgärtner in California (Gutierrez and Baumgärtner 1984). The development of PBDMs flourished under the auspices of the NSF/EPA/USDA funded Huffaker Projects during the 1970s (https://nap.nationalacademies.org/read/9649/chapter/9) enabling deconstruction and analyses of some agricultural systems sufficient to provide sound management recommendations. Two recent holistic PBDM/TVLT analyses of complex systems are the coffee system (Cure *et al*. 2020) and the hybrid Bt cotton system in India (Gutierrez *et al*. 2020, 2024).

Multitude life history strategies occur in nature, but the developmental biology of all multicellular organisms have analogous processes allowing the same modeling paradigm to be used (Gutierrez 1996, Baumgärtner 2020). For example, plants may germinate from seed, grow vegetatively, and mature and produce seed (**Fig. 2a**), and an animal may start as an egg, hatch, and develop as an immature organism passing through various life stages to become a reproducing adult (**Fig. 2b**). Stylized age specific survivorship (*lx*) and reproductive (*mx*) profiles for plants and arthropods are illustrated in the respective lower panels. The life span of the organisms may be hundreds of years (e.g., redwoods) or a few days (e.g., water fleas), but the underling patterns remain the same. Furthermore, the developmental biology of a species may be determinate or indeterminate, and the developmental times of a cohort initiated at the same time have mean and variance. This biology can be captured in PBDMs based on distributed maturation time population dynamics models (see below).

**Figure 2.**
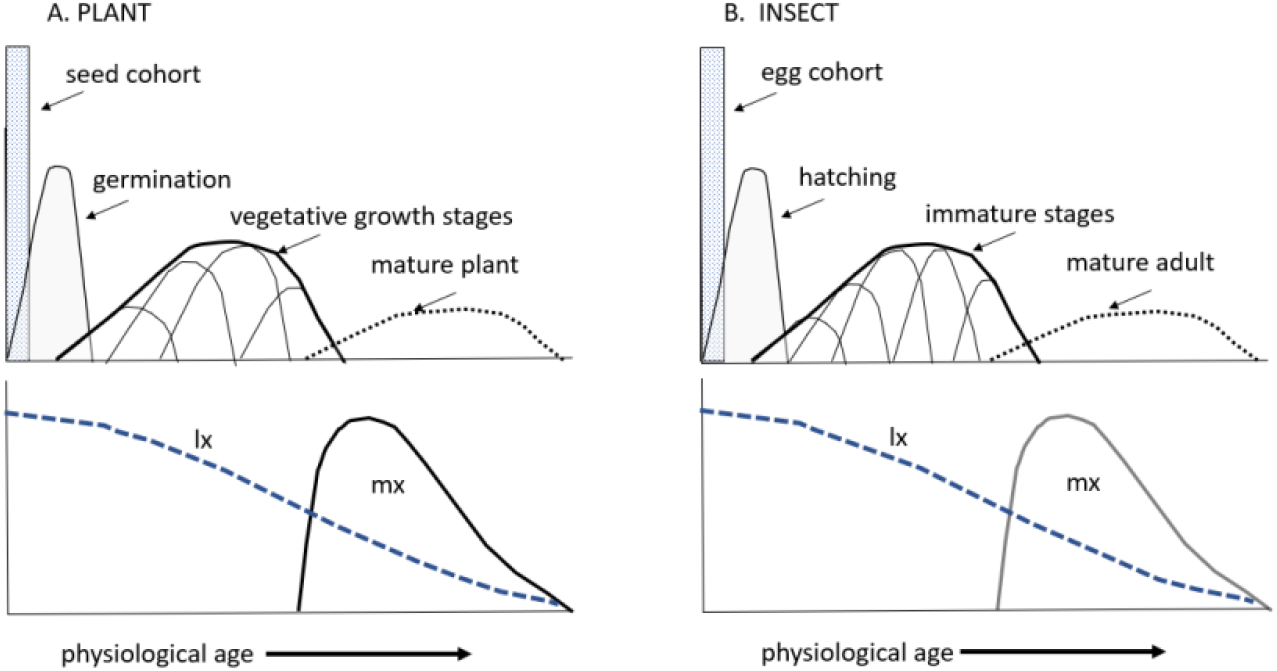
Analogous developmental stages of a plant and an insect with the bottom panel characterizing survivorship (*lx*) and reproductive profiles (*mx*). Not that the developmental time of individuals initiated at the same time have distributed developmental rates.

To capture the field biology of species, age structured PBDM/TVLT (hereafter PBDMs) have been developed using two approaches: (1) the **metabolic pool** (MP) approach of energy acquisition and allocation, and (2) the **biodemographic function** (BDF) approach (Gutierrez 1996). In developing tritrophic system models, all the models may be MP or BDF based, or they may be a mix of the two approaches (e.g., **Fig. 3b,c**). As indicated below, the two approaches are related. The models reported below are the basis for further development.

**Figure 3.**
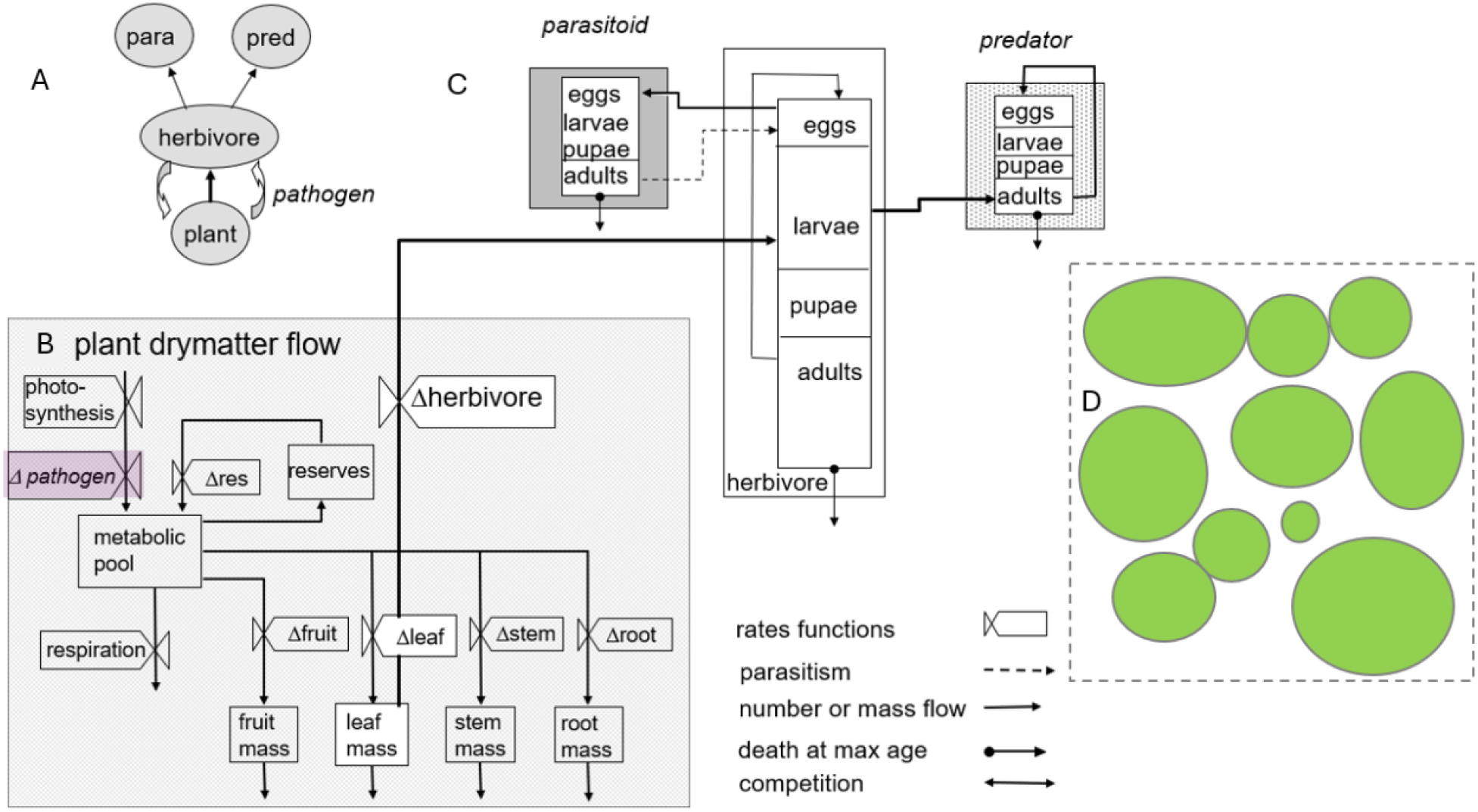
PBDM tri-trophic system: (a) energy/dry matter flow in a tri-trophic system, (b) energy flow within a plant as affected by a pathogen, (c) to higher trophic levels with the herbivore attacking leaves (Gutierrez 1996), and (D) a population of plant each with herbivore/natural enemy populations on them with movement between plants.

### Metabolic pool approach (MP)

MP models are bioeconomic models of how organisms acquire and allocate resources, and by analogy can be used to model the growth and development of animals (insects) (Gutierrez *et al*. 1987), and for harvesting by human firms (Regev *et al*. 1998).

Physiological studies by de Wit and Goudriaan (1978) and colleagues in the Netherlands on plant growth and development provided early impetus for the development of plant models globally. The models sought to capture weather driven processes of resource acquisition and allocation (i.e., photosynthate) for plant growth and development, first to egestion, then to respiration, conversion costs and to growth and reserves if immature, and to reproduction by mature organisms. Gutierrez *et al*. (1975, 1987, see Wang *et al*. 1977) used these notions to develop demographic models of plants composed of age-mass structured populations of leaves, stem, root, and an age-mass-numbers structured population of fruit (**Fig. 3b**; a canopy model). We call these metabolic pool models (**MP**), where the ratio (S/D) of the amount of resource obtained (S, supply) and the physiological demand for resources (D, demand) regulates all vital growth and developmental rates. In PBDM/MP, the demand (D) for photosynthate in plants is the sum of the genetic maximum growth rates of all plant subunits under conditions at time *t* corrected for metabolic costs (e.g., egestion, respiration, and conversion costs). The supply accrued is less than the demand due to imperfect search for resources (e.g., light, water, and nutrients) with the ratio of 0 < 1-S/D < 1 estimating the shortfall and determining the realized growth, death, and reproductive rates (Gutierrez 1992, Gutierrez *et al*. 1994).

Populations of herbivores and natural enemies (parasites, predators, and pathogens, see below) inhabit the canopy, with herbivores attacking specific plant organs in an age specific manner, and natural enemies attack the herbivore in an age specific manner (e.g., Fig. 3C). The dynamics of the system are moderated by plant bottom-up and herbivore/natural enemy top-down effects on species specific S/D ratios (**Fig. 3B,C**).

The canopy model was expanded to a plant population wherein individual cassava plants of different sizes, ages and areas for growth compete with neighbors for light, water, and inorganic nutrients. Furthermore, each plant has interacting populations of herbivores and natural enemies (parasites, predators, and pathogens) on them with species *S/D* influencing movement rates between plants.

### Biodemographic function approach (BDF)

Data to develop MP based PBDMs are often unavailable, and the simpler BDF approach is used. Specifically, birth and death rates of species are estimated from age specific life table studies conducted under different conditions (Gutierrez and Ponti 2013b, Gutierrez *et al*. 2021), and when well done provide a coherent body of knowledge (e.g., Messenger 1964). The predicted age specific life table vital rates are the outcomes over the life cycle of a cohort of organisms of how they acquire and allocate resources, survive, and reproduce under the experimental conditions – i.e., the results of metabolic pool processes. BDFs summarize the developmental, birth and death rates for each species across the experimental conditions and are used to parameterize age-structured, weather-driven PBDMs. The BDFs are summarized in a stylized way in **Figure 4**, though in practice, BDFs fitted to data invariably are not symmetrical. An important shortcoming of this approach is that experiments to characterize effects under extreme conditions are often available – e.g., critical mortality rates (see below).

**Figure 4.**
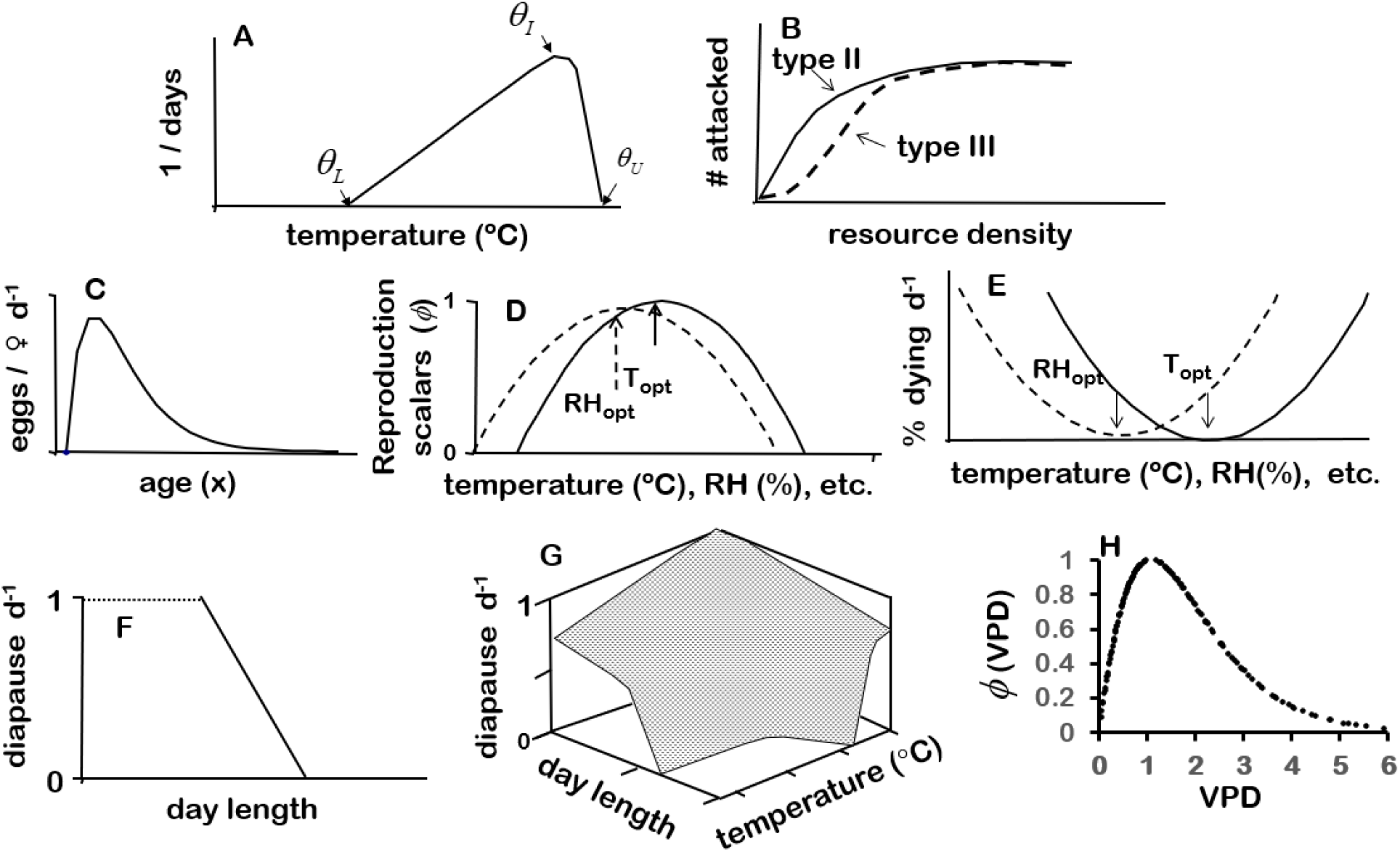
Stylized biodemographic functions (modified from Gutierrez and Ponti 2013a): (a) rate of development on temperature, (B) resource acquisition -the functional response, (C) age specific reproductive profile at the optimum temperature, (D) temperature and relative humidity scalars to correct reproduction from the optimum (E) mortality rate per day at different temperatures and relative humidity, (F, G) the proportion entering dormancy due to photoperiod and due to photoperiod and temperature respectively, and (H) scalar function for oviposition due to vapor pressure deficit.

#### Developmental rates

The developmental rate of poikilotherm plants and animals and their sub stages (*r(T)*, **Fig. 4A**) largely depends on temperature (de Candolle 1855), but other factors such as nutrition may affect *r(T)*. A simple model (eqn. 1) with parameters a and b captures the effects of temperature on *r(T)*, with lower (*θ*_*L*_) and upper (*θ*_*U*_) thresholds for development (*T*, °C) with *θ*_*I*_ being the inflection point where *r(T)* departs strongly from linearity. Other similar functions also suffice (e.g., Briére *et al*. 1999).

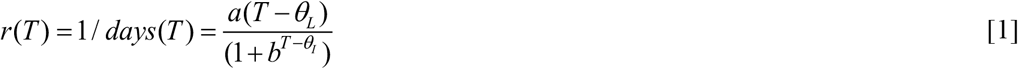

On average, a cohort of individuals initiated at time *t*_*0*_ completes development when 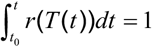. Note that in multi species systems, each (and their stages) may develop on different time scales.

#### Functional response

All organisms are consumers and acquire resources though search (i.e., the functional response, **Fig. 4B**) that may be influenced by several factors. This is also a factor in MP models. A critical innovations for S/D PBDM development was the demand-driven parasitoid ratio dependent Gilbert-Fraser functional model (Frazer and Gilbert 1976; eqn. 2i), and the derived Gutierrez-Baumgärtner predator form (Gutierrez 1996; eqn. 2ii). The parasitoid form is appropriate when a host can be attacked more than once by one or more consumers (e.g., fruit or parasitoid host). In contrast, the predator form posits the resource is consumed and unavailable to other consumers, be it a leaf capturing a quantum of light or a coccinellid beetle consuming an aphid. In both functional response models, *R* is the resource level (be it energy, mass or numbers), *D* is the per capita demand, *α* is the proportion of the available resource that can be found during a time step, and *S* is the amount of the resource acquired by the consumer population *N*. Note that if *α* is a constant, the model is type II (solid line in **Fig. 4B**), but is type III (dashed line) if α is a function of consumer density. The ratio 0 < *S* / *DN* <1 is a metric of resource acquisition success by the population.

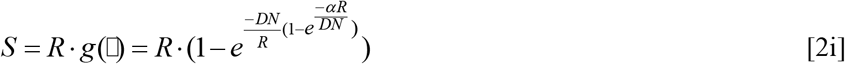

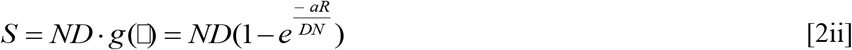

The functional response model becomes more complicated when age structure is added. For example, if adult female parasitoids seek hosts (eqn.2i), the total oviposition site demand by all females in the population 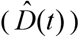 at time *t* is the sum of the product of the age-specific per capita fecundity (i.e., maximum average eggs per female of age (0 ≤ *i* ≤ ^*A*^*k*) per day at optimal temperature *T*_*opt*_, *f* (*i,T*_*opt*_) = *ci/* (1+ *d*^*i*^)) with fitted constants *c* and *d* (**Fig. 4C**), where ^A^*N(i,t)* is the number of adults of age *i* corrected for sex ratio (*sr*) (eqn. 3i).

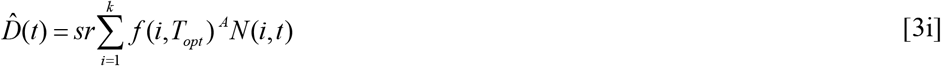

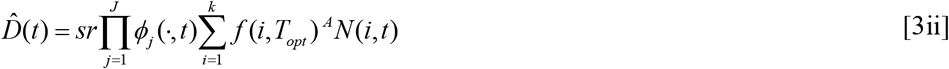

Oviposition demand may be limited by *J* factors, say by temperature (0 < *ϕ*_*T*_ (*t*) ≤ 1) with lower and upper oviposition thresholds, relative humidity (0 < *ϕ*_*RH*_ (*t*) ≤ 1) (**Fig. 4D**), or host age preferences, nutritional effects, and behavior, etc. These limiting factors may be viewed as concave probability functions, and are included in eqn. 3ii as 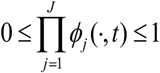 (Gutierrez 1996). In this example, the total population demand 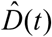 (eqn. 3ii) replaces *DN* in eqn. 2a.

Similarly, the age structured demand of a predator drives its search for any resource be it plant leaves searching for light, roots searching for water and nutrients (e.g., *N, P, K*) or an insect predator searching for prey -all are the same functional processes (**eqn. 2ii**). Detailed physiology or behavior can be added without complication. For example, the mechanistic Ritchie *et al*. (1999) has been used to develop a water balance model to estimate water S/D effects on photosynthesis. Desiccation may also affect orgaism development, survival, and reproduction, and may be included as a scalar of say, oviposition demand (**Fig. 4H**). Vapor pressure deficit (VPD) is a measure of the atmospheric desiccation strength in kPa (kilopascals) (eqn. 4; see Grossiord *et al*. 2020).

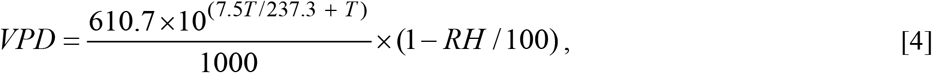

where *T* is the average daily temperature and *RH* is the average daily percent relative humidity. The effects of VPD can be introduced as a normalized concave index scalar (e.g., *ϕ*_*VPD*_ (*t,T, RH*), eqn. 5) in eqn. 3.

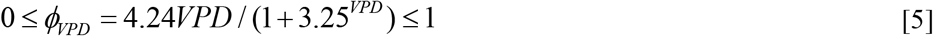

#### Mortality rates

The shapes of the BDF for mortality on temperature and relative humidity illustrated in **Fig. 4E** are symmetrical but the shapes may vary greatly as species may have very different tolerance to temperature extremes and to other factors. Average mortality rates in the favorable range can be estimated as the average slope of the *lx* functions at each temperature in the life table studies (see **Fig. 1**). However, species may experience extreme temperatures, relative humidity or other factors in the field that are unfavorable for development and say the mortality rate may be estimated by exposing cohorts to such conditions. Furthermore, corrections can be made to accommodate daily fluctuating temperature extremes and conditions in protected refuges. The mortality rate (*µ*) is implemented as the product of say the *L* survivorship rates (i.e.,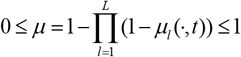), and may be viewed as an applcation of the classic Liebig’s Law of the Minimum (1840).

#### Dormancy

Species may evolve dormancy mechanisms to escape unfavorable periods that may be triggered by temperature, moisture deficits, daylength and other factors. **Figure 4F** shows a simple relationship of dormancy on day length, and **Fig. 4G** shows a more complex relationship on temperature and photoperiod (see Gutierrez *et al*. 1981). Capturing entry to and exit from dormancy may determine how well the model captures the biology determining the geographic distribution and relative abundance and dynamics of a species. Exit from diapause may be viewed as a dynamic stage structured process (Johnsen et al 1997).

### Population dynamics models

Population dynamics models must accommodate the fact that cohort members entering the population at the same time may have different developmental times. Relevant models were reviewed by Gutierrez (1996), Di Cola *et al*. (1999), and Buffoni and Pasquali (2007). For simplicity and ease of implementation, we use the discrete form of the time-invariant and time-varying Erlang distributed-maturation time demographic models (see Abkin and Wolf (1976), Manetsch (1976), Vansickle (1977)) parameterized using MP and/or BDF biology.

Species have different life stages (left superscript s), but the same discrete population dynamics model is used for all of them. The dynamics of a life stage *s* with *k*=1, 2,…,^*s*^*k* age classes (eqn. 6. Manetsch (1976), Gutierrez (1996), Severini *et al*. (2005)) can be viewed as composed of ^s^*k* dynamics equations. The forcing variable is temperature (*T*), with time (*t*) being a day (*d*) that from the perspective of poikilotherm species is of variable length in physiological time units (i.e., 0 < ^s^*Δ*_*x*_*(T(t))* in degree days (*dd*)), or proportional development (^s^*r(T(t))*). Each species (and life stage) may have different mean developmental times (^s^Δ) and temperature thresholds allowing their development on different time scales (see **Fig. 4A**). The biological data used to parameterize PBDMs are most often means collected at daily or longer time intervals, but shorter time steps may be important when daily fluctuation to temperature extremes adversely affect death and other vital rates. The state variable ^*s*^*N*_*i*_*(t)* is the density of the *i*^th^ age class (mass or numbers), and ^*s*^*µ*_*i*_ (*t*) is the proportional age specific net loss rate due to temperature, net immigration, growth in mass dynamics models, and other *L* factors (above) during ^s^*Δ*_*x*_*(T(t))*. Following the notation of Di Cola *et al*. (1999, page 523), the dynamics of the *i*^th^ age class of stage *s* is modeled as follows:

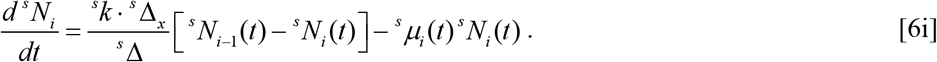

In terms of flux, 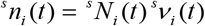 where 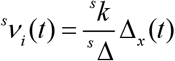, and

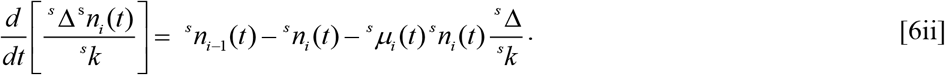

The total density in life stage *s* is 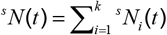.

Ignoring stage notation, new individuals enter the first age class of a stage (*i=*1), flows occur via aging between age classes and between stages at temperature dependent rates, with surviving adults exiting as deaths at maximum age (*i* = ^A^*k*). Absent mortality, the theoretical distribution of cohort developmental times of stage *s* may be estimated by Erlang parameter ^*s*^*k* = ^*s*^ Δ^2^ / ^*s*^*σ* ^2^, where *σ* ^2^ is the variance of average developmental time ^s^Δ. In practice, ^*s*^*k* is an approximation of data. Furthermore, developmental times (rates) may vary with nutrition and other factors (Gutierrez 1996), and given appropriate data can be easily accommodated using the time varying form of the model (Vansickle 1977, see Stone and Gutierrez. I986). Last, because of non-linearities and time varying nature, the model can only be evaluated numerically (Wang and Gutierrez 1980). The numerical solution for eqn. 6ii can be found in Abkin and Wolf (1976), and as implemented here in Gutierrez (1996, pages 157-159).

The algorithm (**Fig. 5**) shows the cycle of the daily computations of birth-death rates and aging for a species (eqn. 6) for a specific location (or for any number of georeferenced lattice cells) across a vast geographic landscape. The flow diagram also illustrates linkages to trophic interactions with other species. The life history variables for all life stages of all species are updated daily in each lattice in runs of several years and the detailed dynamics of system components in any lattice cell may be graphed (e.g., Ponti *et al*. 2021). The output interval (daily, monthly, or yearly) may be specified, and the georeferenced output variables written to time specific text files. In practice, when computing annual average (*x*), standard deviation (std) and coefficient of variation (CV) say across years for all variables in each lattice cell, the first-year results are ignored as the model is assumed equilibrating to site specific conditions. Selected summary variables are mapped using GIS, and marginal analysis (∂*y/*∂*x*_*i*_) may be performed on the full data to estimate the impact of say species or factor *x*_*i*_ on some dependent variable *y* of interest (Gutierrez 1996).

**Figure 5.**
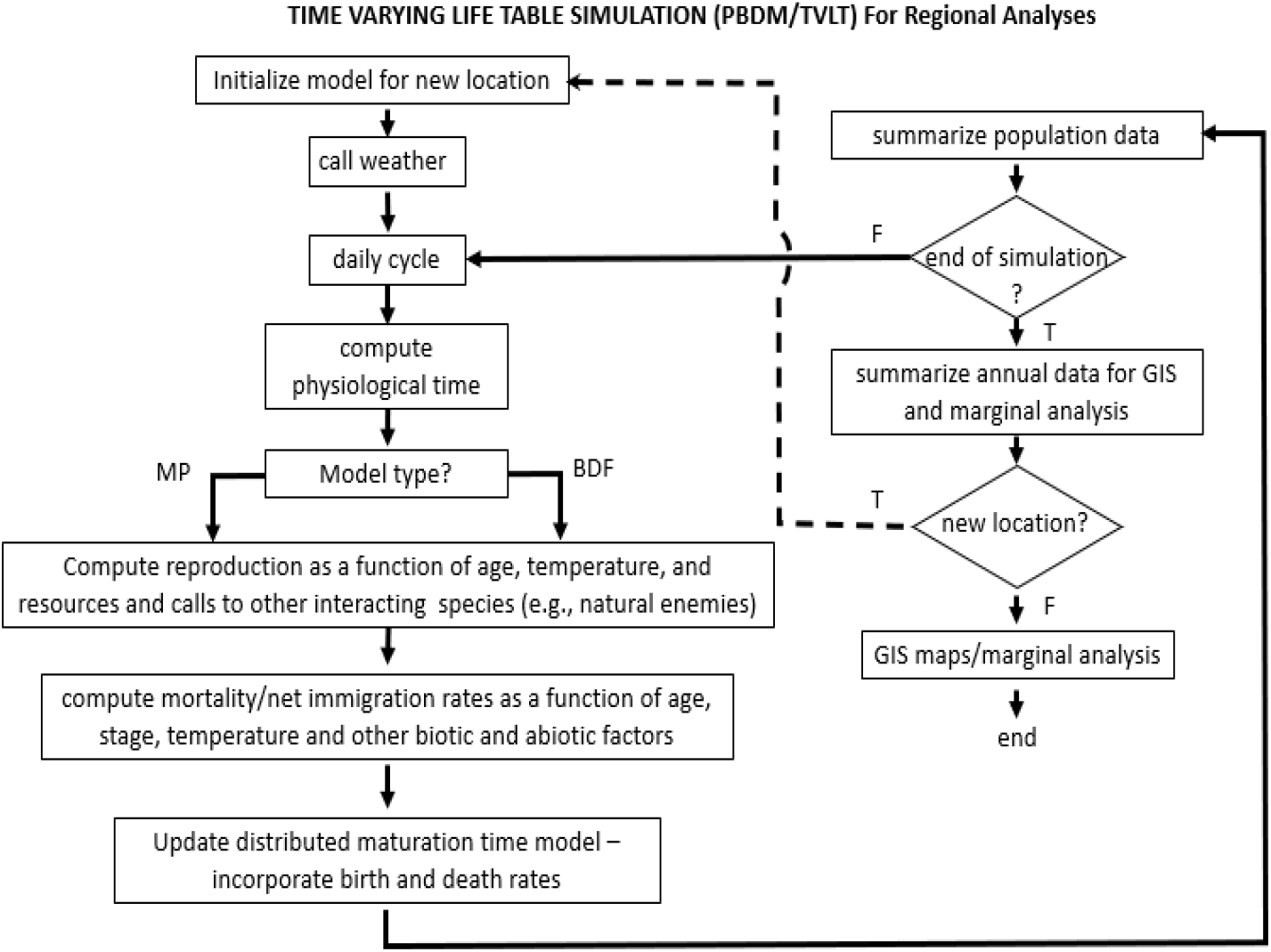
Flow diagram for physiologically based demographic models using metabolic pool (MP) or -biodemographic functions (BDFs) approaches.

### Weather data

The growth and abundance of a species at a geographic location emerges in response to local resource availability, interaction with other species, and weather, all summarized in eqn. 6. From the perspective of an ectotherm species, climate change is simply another weather pattern, and in the extreme determines whether a species can persist and/or invade new areas. The effects of climate change on the systems are not covered due to space constraints, but numerous examples are found in the cited references (see Supplemental Materials Table 1). In this study, the map of Africa was divided into 40,691 georeferenced lattice cells each 25 km^2^, with observed daily weather for each sourced from the global weather dataset AgMERRA (Climate Forcing Dataset for Agricultural Modeling) created as an element of the Agricultural Model Intercomparison and Improvement Project (AgMIP, https://agmip.org/). Because of computation constraints on a laptop computer, a course grid of 10,345 lattice cells was used that capture the relevant details. AgMERRA consists of daily time series over the 1980-2010 period (Ruane *et al*. 2015), and can be accessed through the Goddard Institute for Space Studies (GISS) of the National Aeronautics and Space Administration (NASA, https://data.giss.nasa.gov/impacts/agmipcf/). A maximum of six weather variables are used to run PBDMs (maximum and minimum temperature, precipitation, solar radiation, relative humidity, and wind). Climate model data can be used to study climate change’s effects on species.

### GIS mapping

The open source GIS software GRASS (Neteler *et al*. 2012, GRASS Development Team 2022), see http://grass.osgeo.org/) was used to map PBDM output data. All GIS data layers used in the analysis are available open access, and most were sourced from the public domain repository *Natural Earth* (https://www.naturalearthdata.com/). Inverse distance weighting or bicubic spline interpolation is used in mapping model output as a continuous raster surface. The smoothed spatial patterns reflect not only the site-specific effects of weather on the biology of the species, but also the resolution of the weather grid.

## Results

The goal of our study is to demonstrate the utility of the PBDM approach, hence recommendations for model enhancements are made, noting that all the models can be improved. Some of the models are new and others have been published. Various levels of biological data were available to parameterize the PBDMs with the full extent of the data reported in the references cited.

In Africa, the time and quantity of rainfall is a major factor limiting the geographic distribution and abundance of agricultural and native plants, associated herbivores, and natural enemies, and of medically and veterinary importance species. As background, the average annual precipitation (masked > 2500 mm) during 2001-2010 and associated coefficients of variation as a percent (CV) are mapped in **Figs. 6A, B**. Rainfall varies widely from areas of high rainfall in West-Central Africa to low rainfall in the deserts in the Sahara and the southwestern reaches of Africa with highest CVs occur in the Saharan areas of northern Africa. The major rivers of Africa are indicated as yellow lines in GIS **Fig. 6A** (see inset).

**Figure 6.**
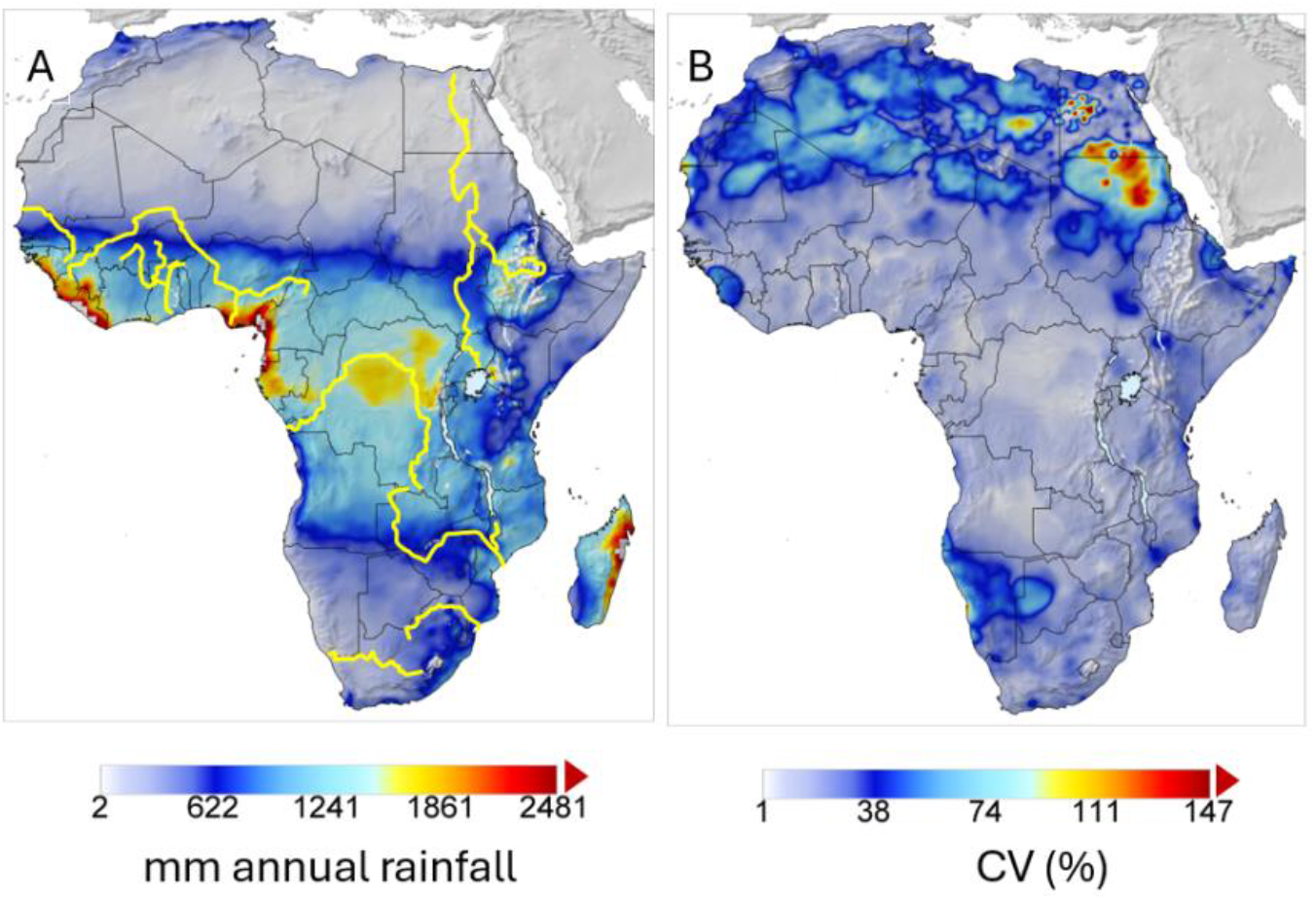
Map of average annual precipitation clipped at 2500 mm (A) and (B) coefficients of variation as a percent for the period 2001-2010 clipped at 150%. The major rivers are indicated as bold yellow lines in A.

### Analysis of species distribution and relative abundance -

#### (1) New-world screwworm

Screwworm (*Cochliomyia hominivorax* (Coquerel) (Diptera: Calliphoridae)) is indigenous to the Western Hemisphere with a wide distribution in subtropical-tropical areas of South, Central and North America and the Caribbean. The fly causes myiasis in warm-blooded animals including humans and hence its egg and larval stages develop at hosts body temperatures (∼38°C internal for cattle). The fly lacks a dormant stage, and the life stages have high lower and upper thermal thresholds for development (**Fig. 7A**; i.e., 14.5^◦^C, 43.5^◦^C respectively), an optimum temperature for egg production of ∼29°C within a range of 14.5–43.5^°^C (see Gutierrez and Ponti 2014a, Gutierrez *et al*. 2019). Adults are sensitive to moderately cold and high temperatures (**Figs. 7B, C**) with lowest mortality occurring at ∼ 27^°^C. In North America, the prospective northern limit of permanence for screwworm is the southern tip of Texas and south Florida characterized by the annual sum of daily mortality rates below the optimum of 27.2°C and less so above it (Gutierrez *et al*., 2019).

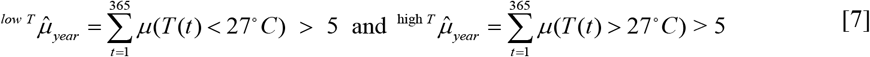

**Figure 7.**
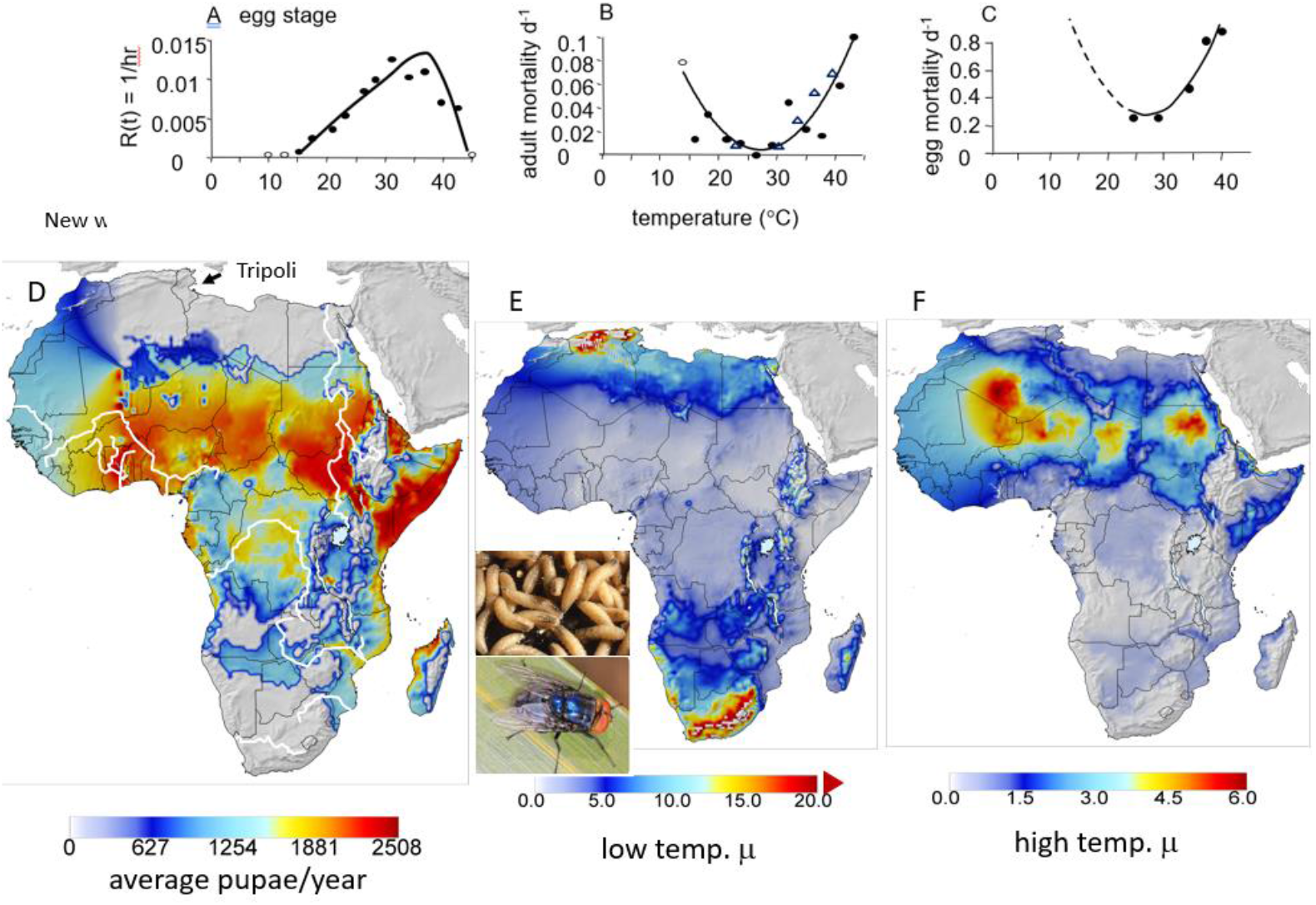
Biodemographic functions and prospective invasiveness of screwworm: (A) Egg developmental rate, (B) adult mortality/day on temperature, (C) and egg mortality rate on temperature, (D) prospective geographic distribution and average relative abundance (i.e., favorability) of screwworm pupae, (E) mean cumulative annual cold weather mortality rate 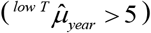 clipped at 20, and (F) average cumulative annual hot weather mortality rate 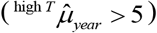. The major rivers in Africa are illustrated as white lines in Fig. 7A.

Historically, periodic outbreaks of myiasis occurred in wildlife and livestock during summer in the southwestern USA (principally Texas) when adult migrants invaded on North American monsoon winds from areas of permanence in northern Mexico (see Gutierrez *et al*. 2019). In the late 1960s, the USDA implemented the sterile insect technique (SIT) (i.e., sterile male) eradication program in the southern USA that was later extended to the Darian Gap in Panama. Screwworm proved to be an ideal candidate for SIT eradication because males are polygynous but a female mate only once, field populations are low, and although the fly has a high reproductive potential, endemic field growth rates are limited by the availability of wound sites for oviposition. Despite an estimated ∼1.25% efficacy rate of released SIT males, eradication in endemic areas of tropical Mexico and Central America was attributable to SIT as the added SIT mating competition on the reproductive success of wild males reduced intrinsically low fly populations leading to native mate-limited extinction (Gutierrez *et al*. 2019).

Screwworm was accidentally introduced to the area of Tripoli, Libya during 1988, and SIT eradication was begun in December 1990 (effectively 1991) (Vargas-Terán *et al*. 2005). PBDM analysis suggests that the area around Tripoli is only moderately favorable for fly permanence, with the vast desert area to the south being unfavorable for the fly due to high temperatures, aridity, and lack of water resources for vertebrate hosts (**Fig. 7D**, Gutierrez and Ponti 2014a). El Azazy (1992) posited that establishment of screwworm along the Nile River (white line) could provide a gateway southward through arid regions of Egypt and Sudan leading to invasion of highly favorable tropical sub-Saharan Africa.

Using the 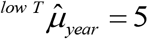(eqn. 7) as a threshold, the favorable areas in Africa are sub-Saharan tropical Africa with decreasing favorability northward and southward due to increasingly cold weather (**Fig. 7E**). Areas of the Sahara are unfavorable due to high temperatures (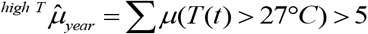 **Fig. 7F**). As a final note, the biological data available to develop the PBDM for screwworm was sparse, this despite approximately one billion US$ spent on screwworm SIT eradication efforts in North and Central America. The comprehensive biology of screwworm needs to be published. The SIT screwworm incidence data for Texas are available on request.

#### (2) Olive/olive fly

Olive (*Olea europaea* L.) is an ancient, drought-tolerant long-lived ubiquitous crop in the Mediterranean Basin that originated in Asia Minor 6000-8000 years ago (https://www.scientificamerican.com/article/the-origins-of-the-olive/). The geographic distribution of olive is limited northward by freezing temperatures and southward by high temperature, and absent irrigation, by low rainfall (Bongi 2002, Vitagliano and Sebastiani 2002, Fiorino 2003). The oligophagous olive fly (*Bactrocera oleae* (Rossi) (Diptera: Tephritidae)) is a major pest of olive, and both species were modeled by Gutierrez *et al*. (2009) and Ponti *et al*. (2009). An important factor is the wider thermal limits in olive relative to olive fly (**Fig. 8A-C**).

**Figure 8.**
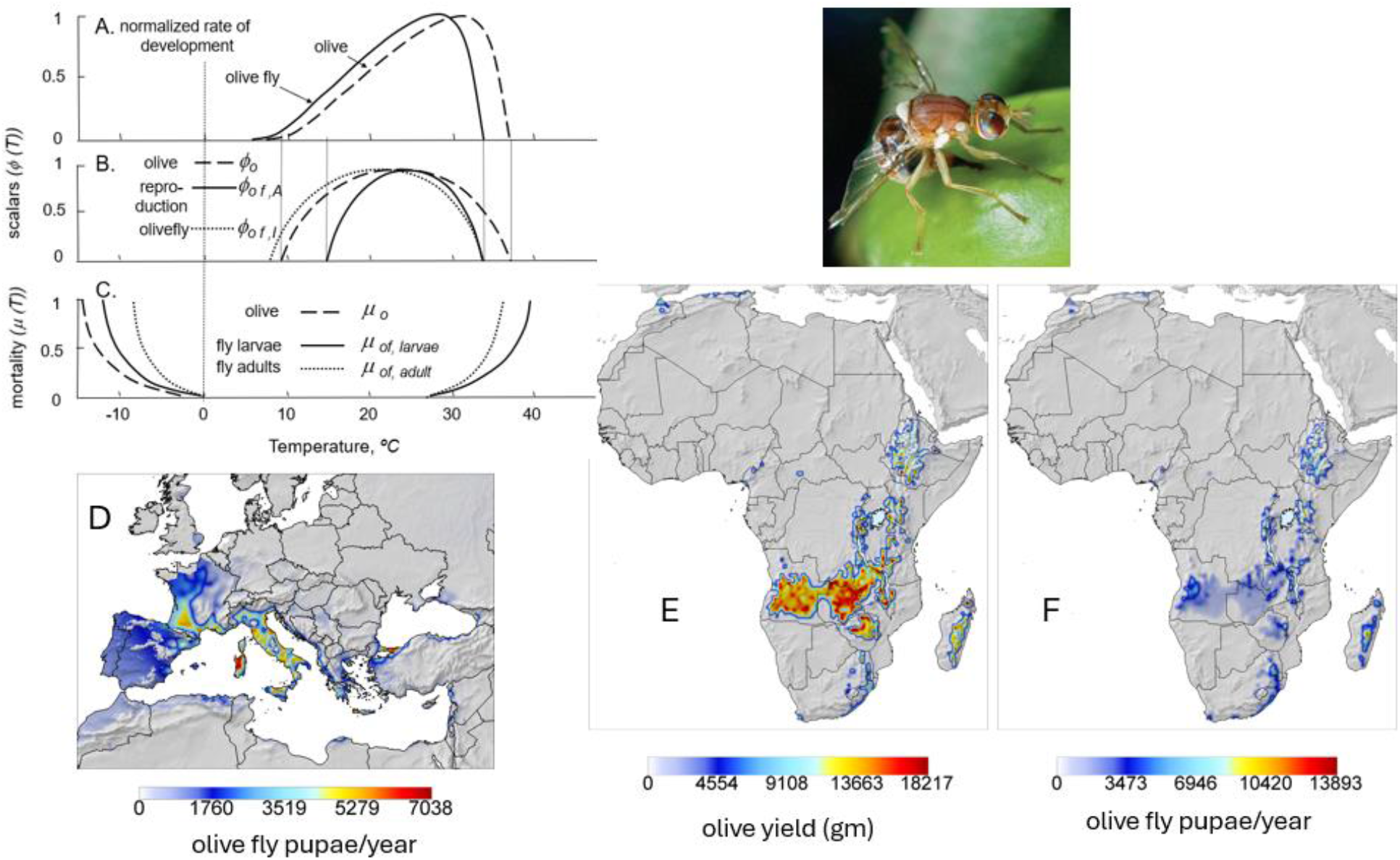
Biodemographic functions for olive and olive fly (A-C, Gutierrez *et al*. 2009), and (D) the prospective distribution olive fly in the Mediterranean Basin, and (D) the prospective distribution olive given annual rainfall >500 mm, and (F) olive fly in Africa. Note that olive is cultivated in Egypt and other regions of North Africa under irrigation.

Relative annual olive yield and cumulative annual olive fly pupae are used as metrics of favorability. For comparative purposes, the simulated prospective geographic distribution and relative abundance of olive fly in the Mediterranean Basin is mapped in **Fig. 8D**. Using the same initial conditions, olive yield and olive fly densities in Africa are mapped in areas above 500 mm annual rainfall in **Figs. 8E,F**. The results show that cooler regions of East Africa and southward into South Africa are favorable for olive, with favorability in areas of Saharan Africa and near coastal areas of North Africa only with sufficient rainfall or irrigation. Olive fly abundance mirrors the distribution of olive, but because the fly is less tolerant to high temperatures, only moderate fly populations are predicted in favorable areas of Africa compared to the Mediterranean Basin (**Fig. 8D vs 8F**).

### (3) Brown marmorated stink bug

The highly destructive polyphagous Asian brown marmorated stinkbug (BMSB), *Halyomorpha halys* (Stål) (Heteroptera: Pentatomidae) invaded North and South America, Europe, and Caucasus region, and here we examine its potential to invade Africa. A tri-trophic PBDM system model of the interactions of BMSB and its parasitoids as forced by weather was used to evaluate prospectively the geographic range of the pest and the impact of natural enemies on its biological control under extant and climate change weather in the Palearctic region (Gutierrez *et al*. 2023). The same model and initial conditions were used to evaluate the potential of BMSB to invade Africa.

BMSB is a temperate species with lower and upper thermal thresholds of 12.1^°^C and 35^°^C respectively, and an optimal temperature for oviposition of ∼26^°^C. The pest is moderately cold tolerant and overwinters as an adult. The analysis includes the effects of two hymenopterous stenophagous egg parasitoids (*Trissolcus japonicus* (Ashmead) and *T. mitsukurii* (Ashmead)), an egg hyperparasitoid *Acroclisoides sinicus* (Huang and Liao), and a tachinid parasitoid flies of the genus (say *Trichopoda* spp.) that attacks large nymphs and adults (**Figs. 9A,B**). The BDFs for BMSB are summarized in **Fig. 9C** and in Gutierrez *et al*. (2023) for the other species.

**Figure 9.**
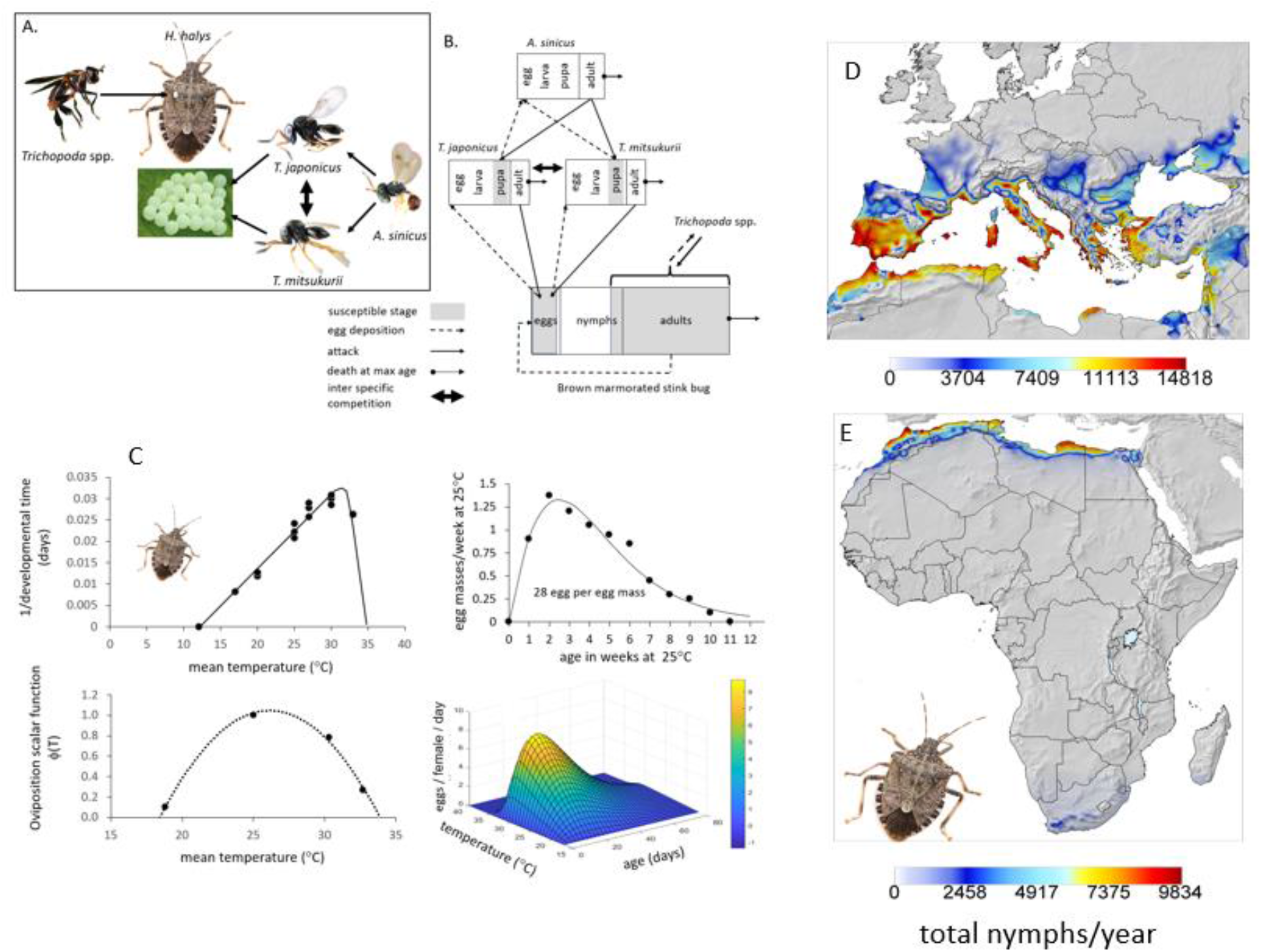
Asian brown marmorated stink bug (BMSB): (A) the trophic relationships, (B) attack relationships, (C) biodemographic functions for BMSB (see Gutierrez *et al*. 2023 for the other species), and the prospective distribution and average relative abundance (i.e., invasiveness) of BMSB nymphs in (D) Palearctic-Mediterranean Region (modified from Gutierrez *et al*. 2023) and (E) Africa.

The prospective range of BMSB is wide throughout temperate areas of Europe, and Mediterranean coastal areas from Morocco across to Egypt (**Fig. 9D**). BMSB is limited by extreme cold temperatures northward in Europe and by hot weather in southern areas. Prospectively, most of the African continent is unfavorable for BMSB with favorability predicted in coastal North Africa and the tip of South Africa (**Fig. 9E**).

### (4) Tomato pinworm

The invasive stenophagous tomato pinworm (*Tuta absoluta* Meyrick (Lepidoptera: Gelechiidae)) is a destructive pest of solanaceous crops. The pinworm is native to the cooler climes of Peru but has extended its geographic range to sub-tropical areas of Brazil. After its initial detection in Spain in 2000, its invasiveness in the Palearctic was analyzed in 2010 and 2019 (see Ponti *et al*. 2021) using the correlative species distribution model CLIMEX developed by Sutherst and Maywald (1985). The CLIMEX studies posited that only coastal southern Europe would be suitable for the pest, despite later records from central Europe, projections were based on its known range expansion in South America into tomato growing areas with warm climates like coastal Mediterranean areas. The analyses failed to consider that the pest originated in the cold semi-arid climes of the Andean highlands, and hence the geographic range of pinworm in Europe proved to be much greater (**Figure 10F**). Its full global invasive potential was not identified until rapid colonization of large areas of Europe had occurred (see Ponti *et al*. 2021), followed by Africa and parts of Asia where it has become a major food security problem.

**Figure 10.**
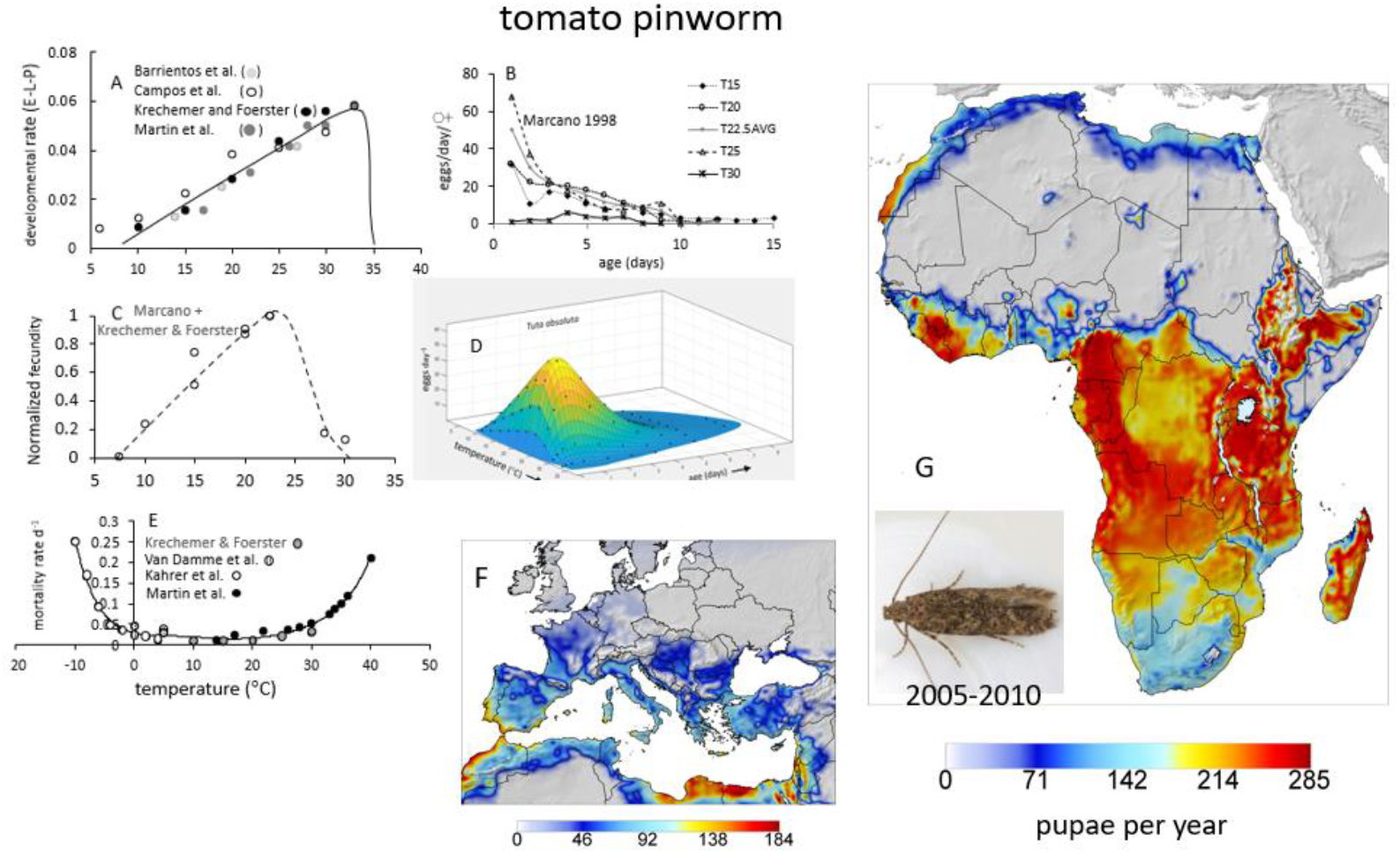
Tomato pinworm: biodemographic functions (A-E), and prospective distribution and average relative abundance in (F) the Palearctic region (see Ponti *et al*. 2021) and (G) in Africa during 2005-2010.

The BDFs for tomato pinworm are illustrated in **Figs. 10A-D**. Its lower thermal threshold is 7.9°C with an upper thermal threshold of ∼35°C, oviposition occurs in the range 7.9 -32°C, the optimal temperature for maximum oviposition is ∼27.5°C, and it has a weak pupal diapause (Campos *et al*. 2021). The mortality rates across temperatures (*µ*_T_*(T)*) were estimated from literature data (Van Damme *et al*. 2015, Krechemer and Foerster 2015, Martins *et al*. 2016, Kahrer *et al*. 2019) (see **Fig. 10A-E**) indicating the moth is tolerant to moderately cold and high temperatures (**Fig. 10E**; see Ponti *et al*. 2021). The excellent biological studies of Campos *et al*. (2021) and Kahrer *et al*. (2019) were critical to this analysis.

The prospective distribution of *T. absoluta* in Africa includes the cooler climes of Senegal, Morocco across coastal North Africa to Egypt and the Levant, with high populations in sub-tropical and tropical Africa, with population levels decreasing in less favorable temperate regions of South Africa (**Fig. 10G**).

### (5) False codling moth

The polyphagous false codling moth (FCM, *Thaumatotibia leucotreta* (Meyrick) (Lepidoptera: Tortricidae)) is native to Africa with an endemic range in Kenya-Ethiopia across to sub-Saharan Africa to West Africa, with an extended range as an invasive species in South Africa with numerous detections in Europe on cut flowers imported from East Africa (EFSA Report 2023). Data on its biology was developed mostly in South Africa (box **Fig. 11A-G**, Daiber 1979a, 1979b, 1979c, 1980, Terblanche *et al*. 2017, and others) and suggest it is adapted to a range of temperatures between 10-35°C with the optimum near 24-25°C. The lower threshold for FCM larval and pupal development is ∼10.5°C with an upper threshold of ∼30°C, females have a short preoviposition period 0.5-1.5 days at 25°C and lay egg singly. Average fecundity increases rapidly above ∼11°C with a sharp decline above ∼26°C to zero at ∼33°C. The moth lacks a diapause stage, and the available temperature dependent mortality data suggest the larval stage is intolerant of freezing temperatures. The larvae are more tolerant to high temperatures than the adult stage (Terblanche *et al*., 2017) captured by a 4°C higher displacement of larval vs. adult mortality rates above 24°C (i.e., the solid line in **Fig. 11G**).

**Figure 11.**
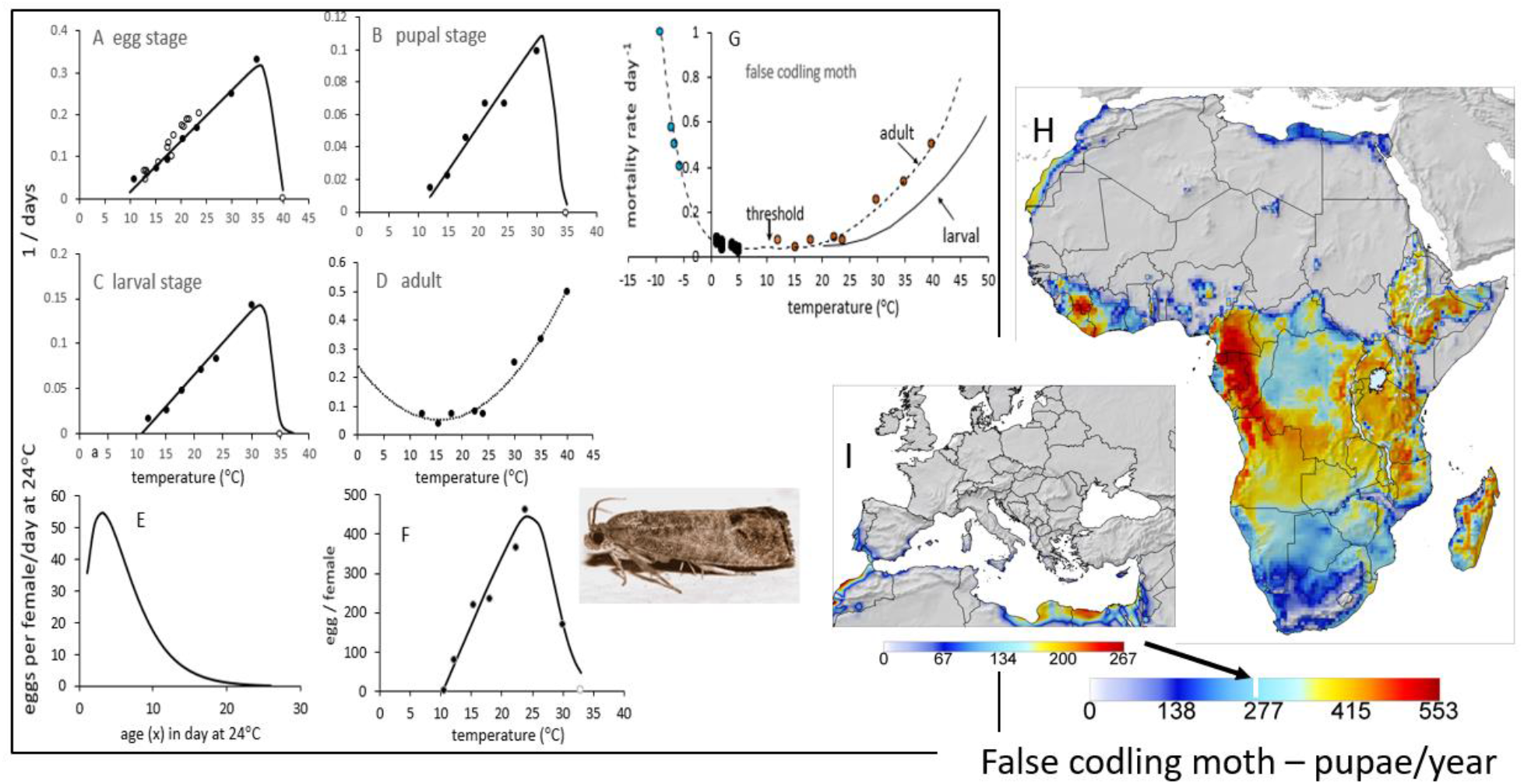
False codling moth: (A-G) biodemographic functions, and prospective distribution and average relative abundance of pupae (H) in Africa and (I) in the Palearctic-Mediterranean Region (see European Food Security Agency 2023).

False codling moth has a documented wide distribution in areas of sub-tropical and tropical Africa (**Fig. 11H**), but its prospective distribution in Mediterranean areas is restricted to the south coastal Iberian Peninsula and north coastal Africa (**Fig. 11I**).

### (6) Fall armyworm

The polyphagous fall armyworm (FAW, *Spodoptera frugiperda* J.E. Smith (Lepidoptera: Noctuidae)) is native to subtropical-tropical regions of the Americas, it has a high migratory capacity, lacks a diapause stage, and has a female biased sex ratio. The moth was first reported in Africa from Nigeria in 2016 (Goergen *et al*. 2016) and is now established across Africa where it causes extensive damage to basic staple crops such as maize, sorghum but also cotton (see Overton *et al*.(2021).

The BDFs for FAW are illustrated in box **Figure 12A-D** showing the larvae and pupae stages have a high lower thermal threshold (12.95°C, 13.25°C respectively) and a relatively high upper thermal threshold of 37.7°C. In the model, the larval threshold is used for the egg and adult stages. At ∼25°C, female moths may produce more than 1,500 eggs over a 14.5-day period, but fecundity declines to zero at 13°C and 37.5°C. The moth is cold intolerant but survives well at high temperatures (**Fig. 12E**), and it is relatively insensitive to changes in relative humidity (see Barfield *et al*. 1978, Ali *et al*. 1990).

**Figure 12.**
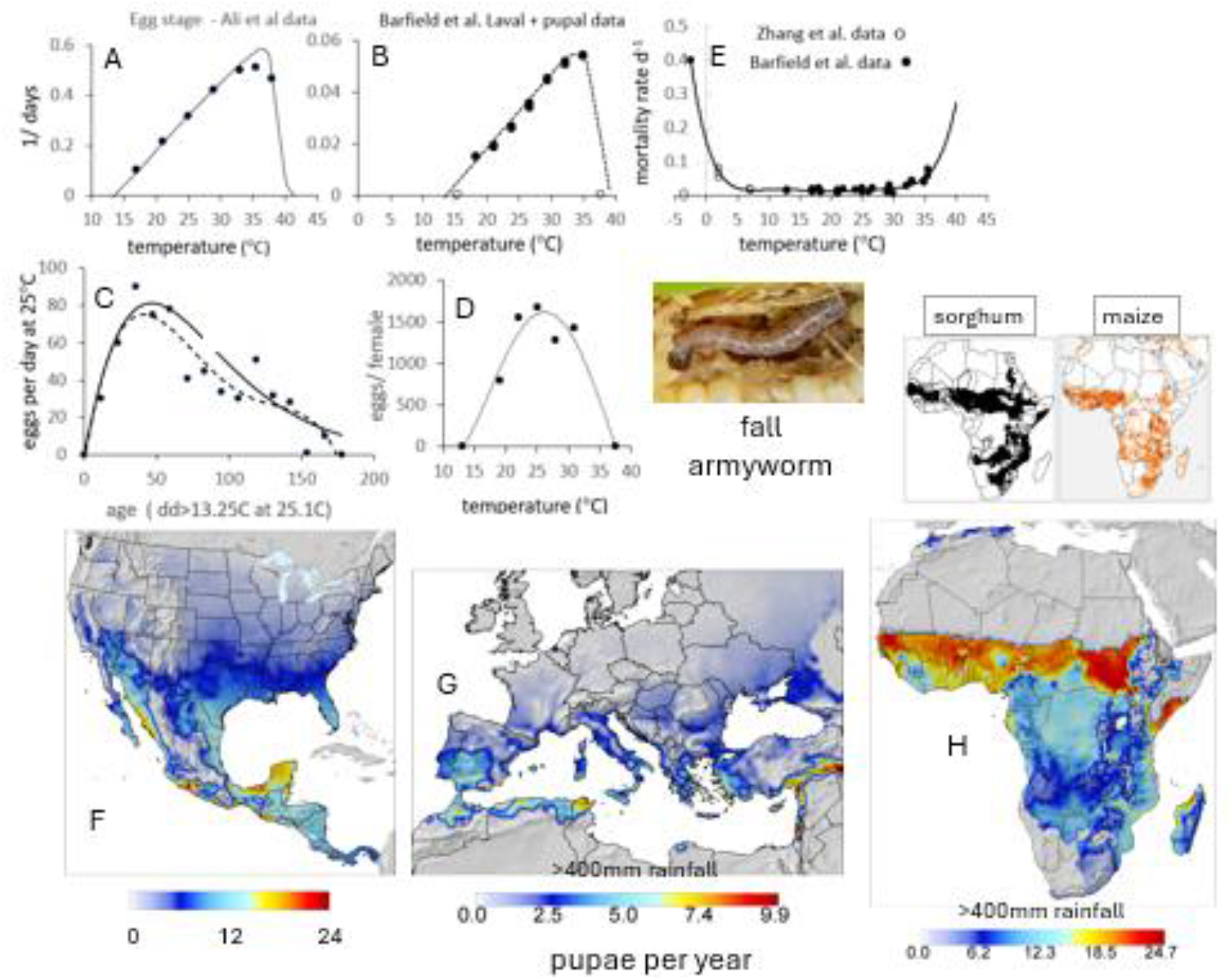
Fall armyworm: **(A-E)** biodemographic functions (e.g., Barfield *et al*. 1978, Ali *et al*. 1990, and others), and using 2000-2010 weather the prospective distribution and average relative abundance of fall armyworm pupae **in (F)** North and Central America, (**G**) the Palearctic-Mediterranean Region and (**H**) Africa. The insets above 12H are the distribution of sorghum and maize (Tang et al. 2024). Map **12H** is masked at >400 mm annual rainfall.

In North America, the prospective permanent distribution of FAW is sub-tropical and tropical areas, with the limits to permanent overwintering being near the midpoint of its prospective density range mapped in **Fig. 12F**. The literature suggests the moth can migrate hundreds of miles on winds that during spring and summer in the USA flow strongly northward carrying moth adults as far as eastern Canada. This allows transient populations to develop in areas of North America where winter temperatures limit year around survival.

Masking areas with less than 400mm annual precipitation, shows that FAW has low capacity to establish in warmer areas of Europe (**Fig. 12G**) confirming the PBDM results of Gilioli *et al*. (2021, 2022). Prospectively, only low populations are predicted in northern Algeria and Tunisia, with higher populations predicted in Egypt and the Levant with prospective maximum density ∼20% lower than in its native range in Mexico and Central America (**12G vs 12H**). The prospective geographic distribution-relative abundance of FAW is a broad band across the continent from Senegal to Somalia-Uganda-Kenya (**Fig. 12H**) with prospective maximum populations about the same as in its native North American range in Mexico (**Fig. 12F**). As is the case in north America, the geographic range of favorability is greater for maize and sorghum than for FAW. The ranges for maize and sorghum in Africa are included above **Fig. 12H** (Tang et al. 2024)

Generalist natural enemies are a strong regulatory component in its native range and needs to be documented in Africa to assess their potential for reducing FAW’s destructive potential. GIS layers for the distribution of its major hosts in Africa would improve the region wide simulations.

#### (7) Citrus/citrus psyllid/parasitoid/greening disease system

The invasive Asian citrus psyllid ((**ACP**), *Diaphorina citri* Kuwayama) (Hemiptera: Liviidae) is a destructive pest causing damage to citrus and to species in 25 genera of Rutaceae (Shivankar *et al*. 2000, Halbert and Manjunath 2004). The psyllid is reported from tropical and subtropical Asia (see Halbert and Manjunath 2004), tropical islands (Reunion, Guadaloupe, Mauritius), parts of South and Central America, Mexico, and the Caribbean (Etienne and Aubert 1980, Shivankar *et al*. 2000, Halbert and Núñez 2004). *D. citri* was first detected in the US in Florida in 1998 (Halbert 1999, Etienne *et al*. 2001), Texas in 2000 (French *et al*. 2001), and southern California in 2008 where concern is that it will spread northward along the coast and into the Great Central Valley (Grafton-Cardwell *et al*. 2013, Grafton-Cardwell and Daugherty 2018). The Asian citrus psyllid has been found in Benin and Nigeria in West Africa and in Kenya, Uganda, and Tanzania in East Africa (Aidoo *et al*. 2020), and its prospective distribution is similar to that of the African citrus psyllid *T. erytreae* (see Aubert 2011, Godefroid 2023).

A PBDM system model was developed to summarizes the available data to assess prospectively the geographic distribution and relative yield of citrus (a generic model), the relative densities of the psyllid, its potential control by the parasitoid (*Tamarixia radiata* Waterston), predation by native predators, and the prospective severity of citrus greening disease in North America and the Mediterranean Basin (Gutierrez and Ponti 2013b). The system of PBDMs is used here to assess the favorability for the species across Africa. The PBDMs for citrus and the psyllid are MP based, and that of the parasitoid is BDF based.

The upper and lower thermal thresholds of the psyllid are 12.85 and 35.5°C respectively with the developmental rate declining to zero above ∼31°C, and the lower thermal threshold for oviposition is about 16°C, the optimum is 31°C and the maximum is about 40°C (box **Fig. 13Aa-Ac**). The psyllid overwinters as an adult but is relatively cold susceptible with the mortality rate d^-1^ being unity at mean -2.5°C (estimated from hourly rates in **Fig. 13Ad**). The upper and lower thermal thresholds *T. radiata* are 7.5 and 35.5°C respectively with the developmental rate declining to zero above ∼31°C, and the lower thermal threshold for oviposition is about 12°C, the optimum is 26°C and the maximum is about 37°C (box **Fig. 13Ba-Bc**). The psyllid’s vital rates are those of a sub-tropical-tropical species, while those of *T. radiata* are more temperate.

**Figure 13.**
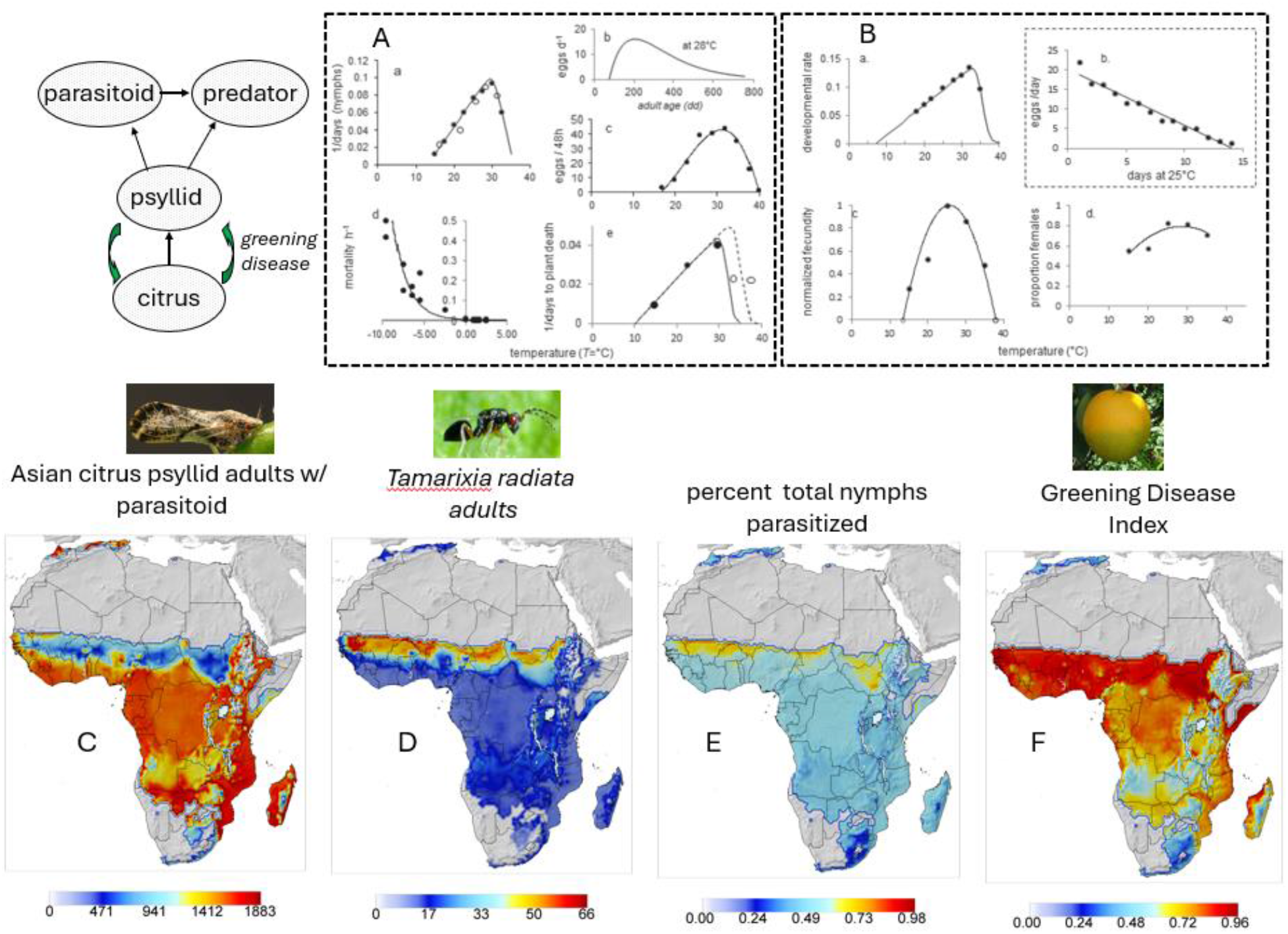
Citrus/Asian citrus psyllid/parasitoid/predator/greening disease system clipped to >400 mm rainfall (see Gutierrez and Ponti 2013b): (**A**) biodemographic functions for ACP on temperature [(**Aa**) developmental rate, (**Ab**) oviposition profile per day at 28°C on age in degree days, (**Ac**) eggs per 48 hours on temperature,(**Ad**) mortality rate per hour on temperature, (**Ae**) rate of infected plant death on temperature, (**B**) biodemographic functions for *T. radiata* on temperature [(**Ba**) developmental rate, (**Bb**) oviposition per day at 25°C, (**Bc**) normalized eggs per day on temperature, (**Bd**) proportion females on temperature, and (**C**) the prospective distribution and average relative abundance of ACP adults/year/tree with parasitism and low predation by indigenous species, (**D**) prospective *T. radiata* adults, (**E**) prospective average percent parasitism of nymphs, and (**F**) greening disease index, (see Aubert 2011).

Absent natural control, the potential distribution of the psyllid is pan-Africa with the highest densities predicted in subtropical areas of southern Africa (not shown). The introduction of the strain of *T. radiata* in our model to Africa would prove ineffective, as psyllid densities (**Fig. 13C**) would prospectively decrease only ∼50% (**Fig. 13D,E**).

ACP is a vector of the bacterial pathogen (‘*Candidatus* Liberibacter asiaticus (CLas)), the causal agent of citrus greening disease (Huanglongbing disease (HLB)) in citrus in many areas of the world, but ACP can also vector two other related phloem-inhabiting fastidious bacteria: *Candidatus* Liberibacter africanus (CLaf) and *Ca*. Liberibacter americanus (CLam). The diseases are transmitted to other trees via clonal propagation, grafting of infected material, with the African citrus psyllid (*Trioza erytreae* (Del Guercio)) being the primary vector of CLaf in Africa (Aubert 2011). Citrus greening disease was recorded in Brazil in 2004 (Coletta-Filho *et al*. 2004), in south Florida in 2005 (Halbert 2005), and in southern California in 2008 (Grafton-Cardwell *et al*. 2014, http://cisr.ucr.edu/Asian_citrus_psyllid.html). Greening disease has wider thermal tolerances than the psyllids, that is reflected in its prospective the geographic distribution of its normalized severity index across tropical areas of Africa is shown in **Fig. 13F**.

A GIS layer for the distribution of citrus and other host Rutaceae in Africa would help define the limits of ACP and HLB. Greater physiological detail for citrus growth and development and greening diseases would improve the model without changing its basic structure. The model can be easily adapted to model the African citrus psyllid.

#### (8, 9) Invasive mosquitoes -*Aedes albopictus* (Skuse, 1894) and *Aedes aegypti* (L.) (Diptera: Culicidae) *Ae*

***albopictus*** is an aggressive human-biting mosquito and is a major vector of important arbovirus diseases including dengue, Zika, and chikungunya. ***Ae. aegypti*** vectors yellow fever, dengue fever, chikungunya, Zika fever, Mayaro and other viral disease agents. A white band on the dorsum of *Ae. albopictus* readily distinguishes it from *Ae. aegypti*.

*Aedes* larvae develop in standing water in natural sites and in artificial containers. The lower thermal thresholds of *Ae. albopictus* and *Ae aegypti* larvae are 10.5°C and 12°C respectively, but the upper inflection points where the developmental rate begins to decline to zero is ∼32.5°C for *Ae. albopictus* and ∼36°C for *Ae. aegypti* (**Figs. 14A, 15A**, Delatte *et al*. 2008, 2009; Bar Zeev 2009a,b; Carrington *et al*. 2013; Brady *et al*. 2013, 2014; Eisen *et al*. 2014; Jia *et al*. 2016). The effects of vapor pressure deficit scalar (0 < *ϕ*_*VPD*_<1) was used to scale oviposition to simulate the effects of dry periods on oviposition by both species. Egg diapause is present in *Ae. albopictus* but not in *Ae. Aegypti* (Perez and Noriega 2013, Denlinger and Armbruster 2014, Diniz *et al*. 2017, Oliva *et al*. 2018) that with it low egg developmental threshold enables it to persist and expand its range in temperate regions (Armbruster 2016). Diapause in *Ae. albopictus* eggs are genetically programmed to environmental change (e.g., photoperiod) which precedes the onset of unfavorable cooler conditions. During diapause, the embryos inside *Ae. albopictus* eggs remains dormant and refractory to environmental stimuli (Denlinger and Armbruster 2014, Jia *et al*. 2016, Batz and Armbruster 2018, Westby and Medley 2020). Quiescence during dry periods occurs in the egg stage of both *Ae. albopictus* and *Ae. aegypti* when the formed embryo (pharate larvae) receives an external stimulus such as a sudden decrease of humidity or a sharp increase in temperature signaling unfavorable conditions for successful hatching. When favorable conditions return, quiescent embryos are sensitive to contact with water that induces rapid hatching (Hand and Podrabsky 2000, Perez and Noriega 2012, 2013, Diniz *et al*. 2017).

**Figure 14.**
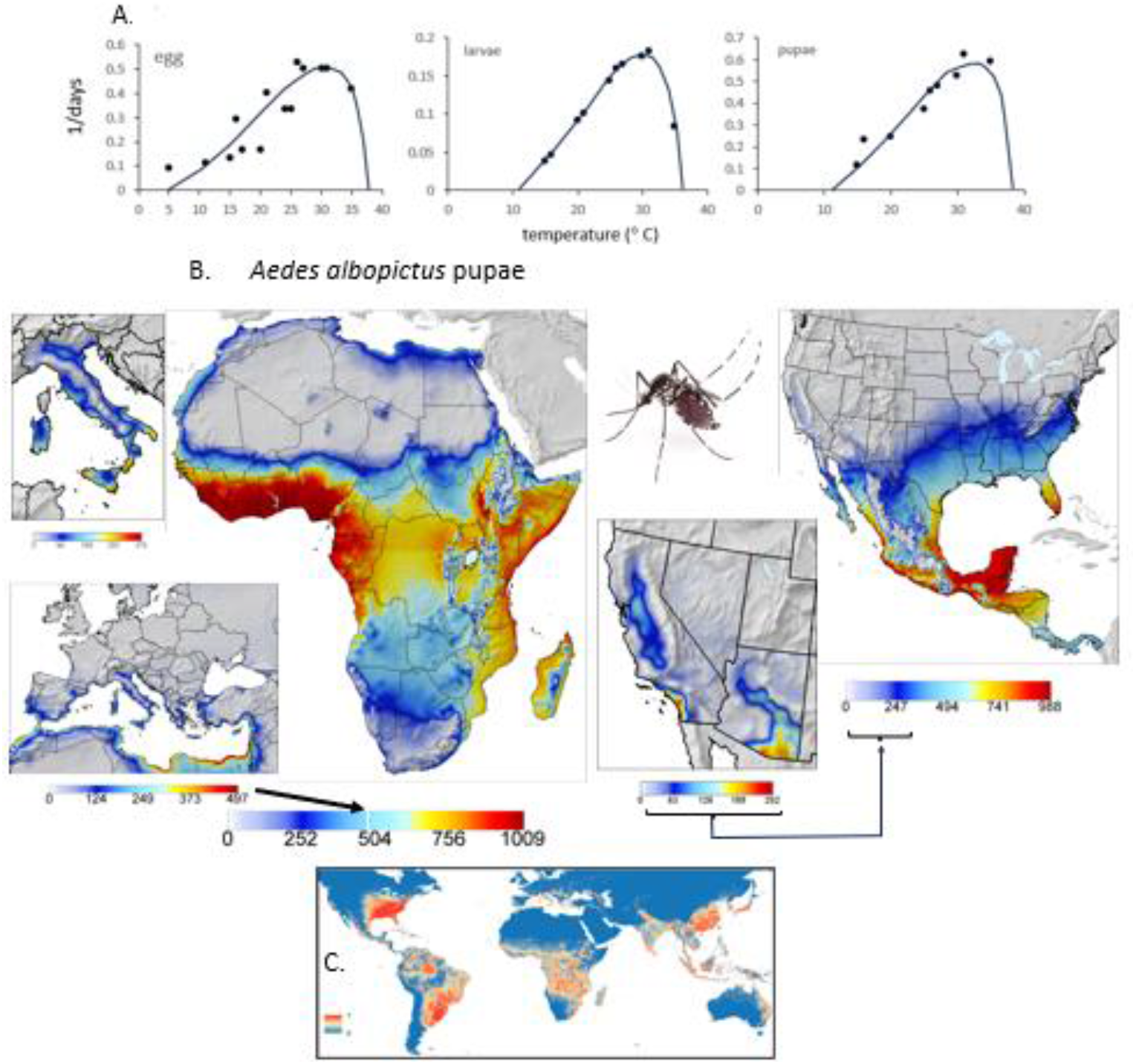
*Aedes albopictus*: (**A**) egg, larval and pupal developmental rates on temperature, (**B**) prospective distribution and average relative abundance in the Palearctic-Mediterranean Region, Africa, and California-Arizona, SDM predictions of mosquito distribution in box (**C**) are from Kraemer *et al*. (2015). The illustration of a female *Ae. albopictus* is from https://www.wechu.org/z-health-topics/aedes-albopictus-mosquito.

The prospective distribution of ***Ae. albopictus*** in temperate regions of the Mediterranean Basin are illustrated in **Fig. 14**B confirming the PBDM predictions of Pasquali *et al*. (2020). Our model predicts the high invasive potential of *Ae. albopictus* in tropical Mexico, lower invasive potential in warmer temperate regions of the SE USA and still lower levels in Coastal southern California and the hot Great Central Valley of California.

Prospectively, ***Ae. aegypti*** does not invade temperate regions of Europe or the Mediterranean Basin (**Fig. 15B**) and compared to *Ae. albopictus* has lower populations levels in Mexico and Central America, and in the hotter regions of California and Arizona.

**Figure 15.**
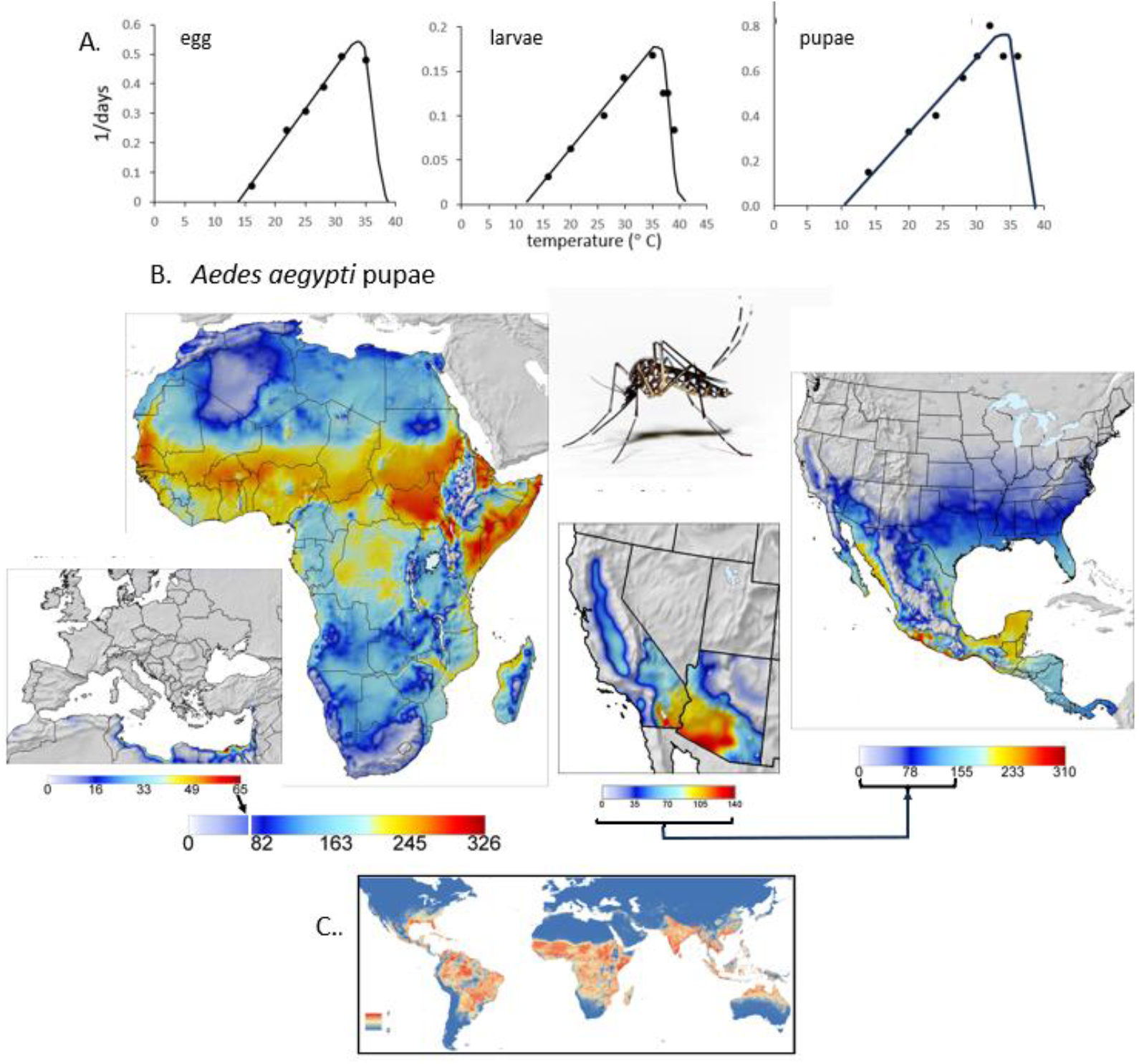
*Aedes aegypti*: (**A**) egg, larval and pupal developmental rates on temperature, (**B**) prospective distribution and average relative abundance of *Aedes aegypti* in the Palearctic-Mediterranean Region, Africa, and California-Arizona. SDM predictions of global mosquito distribution in box C are from Kraemer *et al*. (2015). The illustration of *Ae. aegypti* is from https://www.theatlantic.com/science/archive/2021/06/dengue-mosquitoes-defanged/619161/.

The simulated prospective distributions of the two species across Africa are similar (**Figs. 14B vs 15B**), but the abundance of *Ae. albopictus* is three to four-fold greater than *Ae. aegypti* in favorable regions. This difference is illustrated by the ratio of cumulative annual pupae ((*Ae. albopictus – Ae. aegypti*)/ *Ae. aegypti*) (**Fig. 16**). Also shown in the figure is the limited distribution of both species in arid regions of Africa. The same bias of higher *Ae. albopictus* density also occurs in California and Arizona (**Figs. 14, 15**).

**Figure 16.**
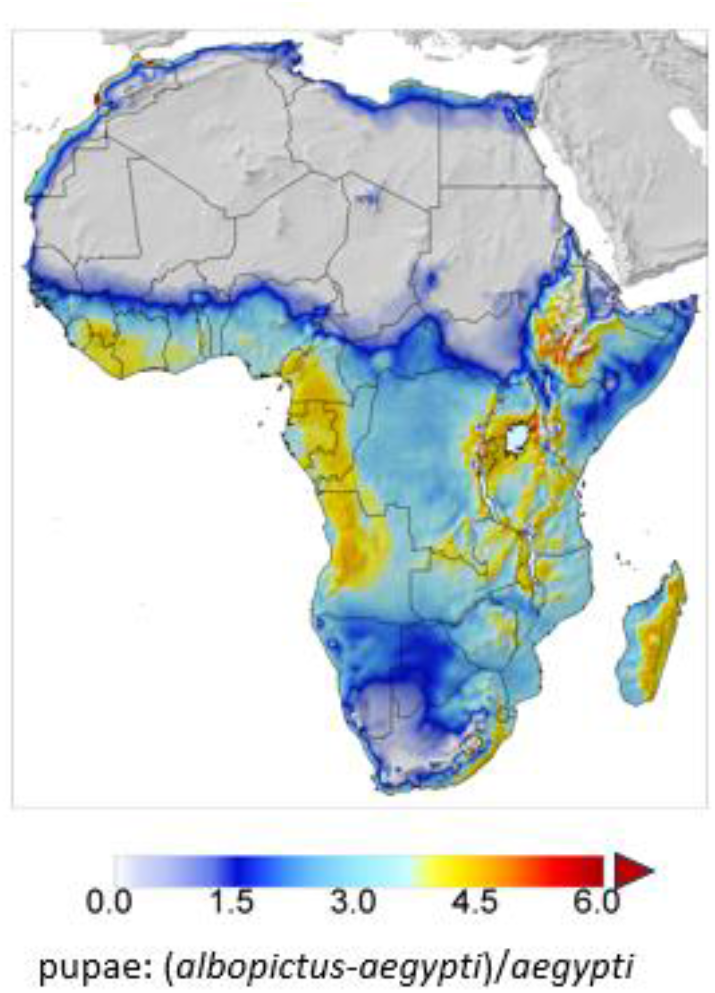
The ratio of pupae ((*Ae. albopictus – Ae. aegypti*)/ *Ae. aegypti*) clipped at 6. The lattice cell values are averages for the ten-year period 2001-2010.

Using presence data and an ensemble Boosted Regression Tree (BRT) approach, Kraemer *et al*. (2015) mapped the global favorability distributions of *Ae. albopictus* and *Ae. aegypti*. They found the distribution of *Ae. albopictus* is sub-tropical and tropical with range extensions into temperate regions (**Fig. 14C**), while the distribution of *Ae. aegypti* is sub-tropical and tropical (**Fig. 15C**). Our results concur with these predictions but were developed independently of species distribution records. As noted earlier, SDM do not capture the biology of the species critical to a holistic understanding of the geographic distribution, dynamics, and relative abundance of species, nor do they allow incorporation of new biological findings on the biology and physiology. For example, biologically rich models are required for regional assessment of population suppression techniques including those based on cytoplasmic incompatibility (Calvitti *et al*. 2010) or the use of *Wolbachia* endosymbionts for control of mosquito populations.

Mosquitoes depend on standing water to breed and a model for standing water dynamic is required, that in its absence the model over predicts favorability in arid areas and during dry periods.

#### (10-13) Four invasive fruit fly species

Sub-tropical and tropical fruit flies are among the most economically important invasive species and are frequently detected by quarantine services in temperate areas. Detection often triggers quarantine and eradication programs conducted without a holistic understanding of the threat posed. A single eradication campaign against tropical fruit flies in the USA costs approximately US$32 million, and up to US$100 million as occurred for medfly in California during 1980-81. Hence, determining the favorability of invaded temperate areas for tropical fruit flies under extant weather and under climate change is critical for policy development.

The most economically important invasive fruit flies are in the family Tephritidae: Diptera from four genera: *Ceratitis, Anastrepha, Bactrocera* and *Rhagoletis*. Tephritid flies have had considerable success in invading new tropical and sub-tropical regions of the world. Their success is attributed to wide host ranges and possibly developmental responses and tolerance to environmental variables through phenotypic plasticity and selections (e.g., Teets and Hahn 2018).

Weather-driven PBDMs of the biology and dynamics of four sub-tropical and tropical species were used to predict prospectively their geographic range and relative abundance (i.e., invasive potential) in North and Central America, and in the European-Mediterranean region under extant and climate change weather (see Gutierrez *et al*. 2021). The four species are: Mediterranean fruit fly (*Ceratitis capitata* (Wiedemann, 1824); medfly) from East Africa, melon fly (*Bactrocera cucurbitae* (Coquillett, 1849)) native to India, Oriental fruit fly (*Bactrocera dorsalis* (Hendel)) from Asia, and the Mexican fruit fly (*Anastrepha ludens* Loew, 1873; mexfly). The detailed biology of these species was reviewed in Gutierrez *et al*. (2021) and not here, though the BDFs are compared in **Fig. 17A-E**. The BDFs of the olive fly (*Bactrocera oleae* (Rossi) (see Fig. 8) and spotted winged fruit fly (*Drosophila suzukii* (Matsumura)(Drosophilidae) that have invaded temperate regions are included in **Figs. 17A-E** to illustrate the relative displacements of their BDFs relative to those of the four sub-tropical-tropical species (Gutierrez *et al*. 2016).

**Figure 17.**
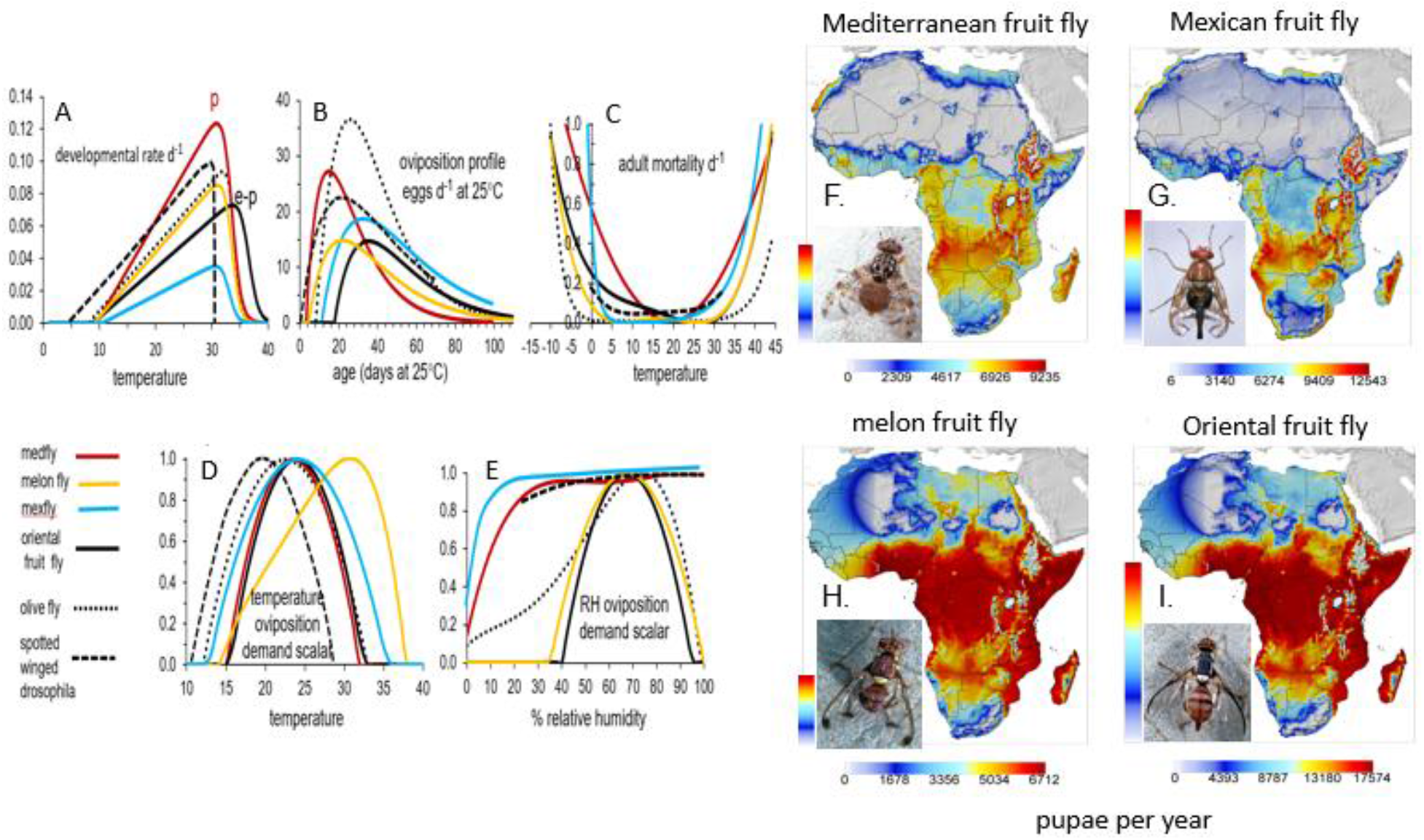
Four invasive fruit flies: (A-E) biodemographic functions (see Gutierrez *et al*. 2021 for BDF details), and prospective distribution and average relative abundance in Africa of (**F**) Mediterranean fruit fly, (**G**) Mexican fruit fly, (**H**) melon fly and (**I**) Oriental fruit fly. (Photos of fruit fly adults are courtesy of Jack Kelly Clark, University of California Statewide IPM Program).

Medfly is native to sub-Saharan Africa and has spread throughout the Mediterranean region, southern Europe, the Middle East, Western Australia, South and Central America, and Hawaii. Mexfly is native to Mexico and Central America, Oriental fruit fly and melon fly are found in tropical regions of the Eastern Hemisphere. None of the four species has a diapause stage and hence cold weather is an important factor restricting their geographic range in temperate regions. Temperature and low relative humidity affect fruit fly reproduction, survival, and permanence in seemingly favorable areas (Gutierrez *et al*. 2021). Prospective distribution maps of the four fruit flies in Africa are **Figs. 17F-I**.

The distribution of Mediterranean fruit fly and Mexican fruit fly across Africa are similar, with the Mexican fruit fly prospectively being ∼25% more abundant in favorable areas (**Fig. 176F, G**). This suggests the Mexican fruit fly could be an important invasive species in sub– Saharan Africa. Melon fly and Oriental fruit fly have similar distributions with Oriental fruit fly being ∼60% more abundant (**Fig. 17H,I**). The vertical color bar in each sub figure are visual comparisons of maximum densities of the four species indicating Oriental fruitfly would be the most abundant.

Improved estimates of the effects of relative humidity on melon fly and Oriental fruit fly are needed and GIS distribution layers of the major host for each of the fruit fly species would enhance mapping their actual geographic range.

## Discussion

Studies of agricultural production are exercises in applied population ecology of crop plants, pests and natural enemies as driven by weather and edaphic, agronomic and economic factors. Further, climate change is simply another weather scenario, and the models readily accommodate this. Most studies on agroecosystems are not holistic, often lack cohesive scientific underpinnings, and become another case histories with limited generality except verbally. What is required to resolve this conundrum is to use analytical methods (models) that capture the underlying biology and dynamics of the species and of the system as driven by weather and resource levels, models that can become libraries where new findings may be added to refine understanding. Mechanistic models of the biology of a species and of the species they interact with address some of these issues.

Physiologically based demographic models (PBDM) bridge some of the gap between field ecology, demography and physiology. They can be developed based on the dynamics of resource acquisition and allocation (i.e., metabolic pool (MP)) or the simpler biodemographic function (BDF) approach that summarizes the vital rates from age specific life table studies conducted under an array of temperatures, relative humidity, levels of nutrition and other factors as appropriate. Age-specific life table statistics are the results of energy/resource acquisition and allocation processes under the experimental conditions – of MP processes. Similar BDFs can be used to describe the weather and resource driven biology of poikilothermic species – of plant and insect species. Data on the biology of species based on MP or BDF approaches were used here to parameterize weather and resource driven distributed maturation time dynamics models. However, no model can be a one-to-one description of the biology, but rather the goal is to capture the key features sufficient to yield sound predictions about the time-place varying distribution, relative abundance, and dynamics of species, and in a GIS context across thousands of lattice locations of vast geographic regions such as Africa – independent of species presence records. PBDMs easily accommodate additional biology and physiology as required without changing the basic character or structure of the models. Detailed daily age structured dynamics of the species are computed and can be examined for any lattice cell location. Our goal was to demonstrate the relative eases of developing PBDMs and demonstrating part of their utility by examining the geographic distribution and relative abundance of thirteen invasive species across Africa. All the models in this study can be updated with additional data and attributes.

Members of the CasasGlobal.org consortium using in-house heritage software have developed and implemented numerous PBDMs in a GIS context (Supplemental Materials **Table 1)** using variations of the underlying MP-BDF PBDM paradigm (see also Gutierrez and Ponti 2013a). However, transferring this technology to others researchers has proven difficult with the major limiting factors training using the software and software incompatibility. To circumvent these apparent barriers for analyzing pest systems rapidly and to assist development of management and implementation policy, a general PBDM/GIS application platform interface (API) is being developed that would enable non-experts to construct PBDMs for various purposes such as examining seemly complex weather driven phenology and dynamic of species and of species interactions at local, regional, and larger scales (Supplemental Materials **Figure 1**).

As a caveat, no model, including PBDMs, can predict crop yield or pest organism densities and damage with a sufficient degree of accuracy for economic speculation. Rather, PBDMs are heuristic tools designed to guide scientific understanding for the development of management practices and policy. Two recent holistic analyses are the Indian hybrid Bt cotton (Gutierrez *et al*. 2020) and Colombian coffee (Cure *et al*. 2020) sysems that explained important biological components of the system and provided management recommendations. Furthermore, PBDMs can be used as the objective function in economic analyses to provide important insights (Regev *et al*. 1998, Pemsl et al. 2007, Gutierrez *et al*. 2020) not captured by econometric analyses of field panel data, and to examine ecological theory (Schreiber and Gutierrez 1998).

## Acknowledgements

We continue to be grateful to the international network maintaining the Geographic Resources Analysis Support System (GRASS) software and making it available to the scientific community. The study was supported by CASAS Global NGO (https://www.casasglobal.org), the McKnight Foundation (grant number 22-341), and project TEBAKA (project ID: ARS01_00815) co-funded by the European Union -ERDF and ESF, “PON Ricerca e Innovazione 2014-2020”. Access to the library search engine of the University of California at Berkeley was invaluable.

