## Supplementary Table 1 and Figure 1 for "Physiologically based demographic model/GIS analyses of thirteen invasive species in Africa: why the biology matters"

[3]Agenzia Nazionale per le Nuove Tecnologie, l'Energia e lo Sviluppo Economico Sostenibile (ENEA), Centro Ricerche Casaccia, Via Anguillarese 301, 00123 Roma, Italy.

[4] mundialis GmbH & Co. KG, Koelnstrasse 99, 53111 Bonn, Germany;

[5]Facultad de Ciencias Básicas y Aplicadas, Universidad Militar Nueva Granada, Cr. No. 101-17 80, Bogotá, Colombia.

[6]IDEMS International, Reading, Berkshire, United Kingdom;

ORCID APG: 0000-0001-7773-1715

ORCID LP: 0000-0003-4972-8265

ORCID JRC: 0000-0001-5816- 0259

ORCID MN: 0000-0003-1916-1966

OCID GS: 0000-0002-9076-4973

**Supplemental Materials: Table 1.** Partial list of published PBDM system models and references with indication of host plant, herbivore species and the number of natural enemy species modeled, geographic location of the study, if in a geographic information system (GIS) context (+ true; - false), and if climate change effects were included (+, -) (see also Gutierrez and Ponti 2014b, Gutierrez *et al.* 2021). References to theory are in the list.

| ***Common name*** | ***Scientific name*** | ***Enemy models*** [a] | ***Geographic location*** | ***GIS*** | ***Climate change*** | ***Plant Host*** |
|---|---|---|---|---|---|---|
| **alfalfa** | *Medicago sativa* | 4 | Western USA | + | + | alfalfa |
| alfalfa weevil | *Hypera postica* | 0 | California, Spain, Iran | + | - | alfalfa |

| *Common name* | *Scientific name* | *Enemy models* [a] | *Geographic location* | *GIS* | *Climate change* | *Plant Host* |
|---|---|---|---|---|---|---|
| blue aphid | *Acyrthosiphon kondoi* | 4 | California | + | + | alfalfa |
| pea aphid | *Acyrthosiphon pisum* | 4 | California | + | + | alfalfa |
| spotted alfalfa aphid | *Therioaphis maculata* | 4 | California | + | + | alfalfa |
| **apple** | *Malus domestica* | 7 | Switzerland | - | - | apple |
| European red spider mite | *Panonychus ulmi* | 0 | Switzerland | - | - | apple |
| two-spotted spider mite | *Tetranychus urticae* | 0 | Switzerland | - | - | apple |
| rosy apple aphid | *Dysaphis plantaginea* | 0 | Switzerland | - | - | apple |
| apple-grass aphid | *Rhopalosiphum insertum* | 0 | Switzerland | - | - | apple |
| apple aphid | *Aphis pomi* | 0 | Switzerland | - | - | apple |
| **common bean** | *Phaseolus vulgaris* | 0 | Colombia | - | - | bean |
| **cassava** | *Manihot esculenta* | 2 | sub–Saharan Africa | - | - | cassava |
| cassava green mite | *Mononychellus tanajoa* | 4 | sub–Saharan Africa | - | - | cassava |
| cassava mealybug | *Phenacoccus manihoti* | 4 | sub–Saharan Africa | - | - | cassava |
| **citrus** | *Citrus* | 1 | USA / Mexico | + | + | citrus |
| Asian Citrus psyllid [c] | *Diaphorina citri* | 2 | USA/Mexico | + | + | citrus |
| **coffee** | *Coffea arabica* | 1 | Brazil-Colombia | + | - | coffee |
| coffee berry borer | *Hypothenemus hampei* | 5 | Brazil-Colombia | - | - | coffee |
| **cowpea** | *Vigna unguiculata* | 0 | Benin | - | - | cowpea |
| bean flower thrips | *Magalurothrips sjoestedti* | 0 | Benin | - | - | cowpea |
| **cotton** [b] | *Gossypium hirsutum* | 7 | global | + | + | cotton |
| beet armyworm | *Spodoptera exigua* | 0 | California | - | + | cotton |
| bollweevil | *Anthonomus grandis* | 0 | California | - | + | cotton |
| cabbage looper | *Trichoplusia ni* | 0 | California | - | + | cotton |
| cotton bollworm | *Helicoverpa zea* | 0 | California | - | + | cotton |
| lygus bug | *Lygus hesperus* | 0 | California | - | + | cotton |
| pink bollworm | *Pectinophora gossypiella* | 0 | Americas, Egypt, India | + | + | cotton |
| white fly | *Bemisia tabaci* | 3 | California | - | - | cotton |
| verticillium wilt | *Verticillium dahliae* | 0 | California | - | - | cotton |
| **grape** | *Vitis vinifera* | 0 | USA / Switzerland / Palearctic | + | + | grape |
| European grape leafhopper | *Scaphoideus titanus* | 2 | Switzerland / Italy | - | - | grape |

| *Common name* | *Scientific name* | *Enemy models [a]* | *Geographic location* | *GIS* | *Climate change* | *Plant Host* |
|---|---|---|---|---|---|---|
| glassy-winged sharpshooter [c] | *Homalodisca vitripennis* | 2 | USA | + | + | grape |
| grape berry moth | *Lobesia botrana* | 0 | USA/Palearctic | + | + | grape |
| European red mite | *Panonychus ulmi* | *0* | Switzerland | - | - | apple |
| grapevine mealybug | *Planococcus ficus* | 3 | California | + | - | grape |
| **olive** | *Olea europaea* | 2 | USA / Palearctic | + | + | olive |
| olive fly | *Bactrocera oleae* | 0 | USA / Palearctic/Africa | + | + | olive |
| olive scale | *Parlatoria oleae* | 2 | California | - | - | olive |
| **rice** | *Oryza sativa* | 0 | Philippines / Madagascar | - | - | rice |
| African white rice borer | *Maliarpha separatella* | 0 | Madagascar | - | - | rice |
| rice aquatic system | *aquatic food web* | food web | Philippines | - | - | rice |
| **yellow starthistle (YST)** | *Centaurea solstitialis* | 5 | USA / Palearctic | + | + | starthistle |
| seed feeder diptera | *Chaetorellia australis* | 0 | USA / Palearctic | + | + | starthistle |
| seed feeder diptera | *Urophora sirunaseva* | 0 | USA / Palearctic | + | + | starthistle |
| seed coleoptera | *Bangasternus orientalis* | 0 | USA / Palearctic | + | + | starthistle |
| YST seed feeding - | *Eustenopus villosus* | 0 | USA / Palearctic | + | + | starthistle |
| YST rosette weevil | *Ceratapion basicorne* | 0 | USA / Palearctic | + | + | starthistle |
| **Individual Species** [d]**:** | | | | | | |
| brown marmorated stink bug | *Halyomorpha halys* | 3 | North America/ Palearctic/Africa | + | + | — |
| bumble bee | *Bombus atratus* | 0 | Colombia | - | - | — |
| cabbage aphid | *Brevicoryne brassicae* | 0 | Australia | - | - | — |
| cabbage rootfly | *Delia radicum* | 0 | California | - | - | — |
| coccinellid beetle | *Hippodamia convergens* | 0 | California | - | - | — |
| cowpea aphid | *Aphis craccivora* | 1 | Australia | - | - | — |
| false codling moth | *Thaumatotibia leucotreta* | 0 | Africa / Palearctic | + | + | — |
| medfly | *Ceratitis capitata* | 0 | North America / Palearctic | + | + | — |
| melon fly | *Bactrocera cucurbitae* | 0 | North America / Palearctic/Africa | + | + | — |
| Mexican fruit fly | *Anastrepha ludens* | 0 | North America / Palearctic/Africa | + | + | — |
| North American screwworm | *Cochliomyia hominivorax* | 0 | North-Central America / Palearctic | + | + | — |

| ***Common name*** | ***Scientific name*** | ***Enemy models[a]*** | ***Geographic location*** | ***GIS*** | ***Climate change*** | ***Plant Host*** |
|---|---|---|---|---|---|---|
| Oriental fruit fly | *Bactrocera dorsalis* | 0 | North America / Palearctic/Africa | + | + | — |
| sheep blowfly | *Lucilia cuprina* | 0 | laboratory | - | - | — |
| New world screwworm | *Cochliomyia hominivorax* | 0 | North America/Africa | + | + | — |
| spotted wing drosophila | *Drosophila suzukii* | 0 | North America/Palearctic | + | + | — |
| tiger mosquito | *Aedes albopictus* | 0 | North America / Palearctic/Africa | + | + | — |
| yellow fever mosquito | *Aedes aegypti* | 0 | North America / Palearctic/Africa | + | + | — |
| thimble berry aphid | *Masonaphis maxima* | 2 | British Columbia | - | - | — |
| tomato pinworm | *Tuta absoluta* | 0 | USA / Palearctic | + | + | — |

[a] The number of natural enemy species included in the study.
[b] incorporation of Bt toxins.
[c] transmission of disease.
[d] plant/animal host not modeled.

**Partial list of references for Supplemental Materials Table 1**

**Asian citrus psyllid**

Gutierrez, A.P., and L. Ponti (2013) Prospective analysis of the geographic distribution and relative abundance of **Asian citrus psyllid** (Hemiptera: Liviidae) and citrus greening disease in North America and the Mediterranean Basin. *Florida Entomologist* 96(4) 1374- 1391. http://doi.org/10.1653/024.096.0417 [Open Access]

**Alfalfa**

Gutierrez, A. P., J. B. Christensen, C. M. Merritt, W. B. Loew, C. G. Summers, and W. R. Cothran. 1976. Alfalfa and the **Egyptian alfalfa weevil**. Can. Ent. 108: 635-648.

Gutierrez, A. P., J. U. Baumgärtner, and K. S. Hagen. 1981. A conceptual model for growth, development, and reproduction in the **ladybird beetle**, *Hippodamia convergens* (Coleoptera: Coccinellidae). Can. Entomol. 113: 21-33.

Gutierrez A.P., Ponti L., 2013. Deconstructing the control of the **spotted alfalfa aphid** (*Therioaphis maculata*). *Agricultural and Forest Entomology*, http://dx.doi.org/10.1111/afe.12015

Gutierrez, A. P., and J. U. Baumgärtner. 1984. Multitrophic level models of predator-prey energetics: I. Age specific energetics models--**pea aphid** *Acyrthosiphon pisum* (Harris) (Homoptera: Aphididae) as an example. Can. Entomol. 116: 923-932.

Gutierrez, A. P. and J. U. Baumgärtner. 1984. Multitrophic level models of predator-prey energetics: II. A realistic model of plant-herbivore- parasitoid-predator interactions. Can. Entomol. 116: 933-949.

Gutierrez, A. P., J. U. Baumgärtner and C. G. Summers. 1984. Multitrophic level models of predator-prey energetics: III. A case study in an alfalfa ecosystem. Can. Entomol. 116: 950-963.

Gutierrez, A.P., L. Ponti, A. Levi-Mourao, X. Pons, J.R. Cure, M. Neteler, G Simmons (in press) Centripetal biotype selection in the invasive alfalfa weevil (*Hypera postica* (Gyllenhal)). J. Applied Ecology

**Apple**

Baumgärtner, J. U., M. Genini, B. Graf, A. P. Gutierrez, and P. Zahner. 1988. Generalizing a population model for simulating Golden Delicious apple tree growth and development. (Book chapter). Proc. Intern. Symp. Computer Modelling in Fruit Research and Orchard Management. Stuttgard-Holenheim

Baumgärtner, J. U., A. P. Gutierrez, and A. Klay. 1988 Elements of modeling the dynamics of tritrophic population interactions. Exp. Appl. Acarology 5: 243-263.

**Bean**

Gutierrez, A.P., E. Mariot , J.R. Hakim Cure and A. Villacorta. 1993. A Model of Bean *(Phaseolus vulgaris* L.) Growth Types I-III Factors Affecting Yield. Agric. Syst. 44:35 - 63.

**Brown marmorated stinkbug**

Gutierrez, Andrew Paul, Giuseppino Sabbatini Peverieri, Luigi Ponti, Lucrezia Giovannini, Pio Federico Roversi, Alberto Mele, Alberto Pozzebon, Davide Scaccini and Kim A Hoelmer (2022) Regional tritrophic analysis of the prospective biological control of **brown marmorated stinkbug** under extant and climate change weather. Pest Science https://doi.org/10.1007/s10340-023-01610-y

**Cabbage root fly**

Johnsen, S., A. P. Gutierrez, and J. Freuler. 1990. The within season population dynamics of the **cabbage root fly** (*Delia radicum* (L.)). A simulation model. Bulletin de la Societe Entomologique Suisse 63 : 451-463.

Johnsen, S; Gutierrez, A P. 1997. Induction and termination of winter diapause in a Californian strain of the **cabbage maggot** (Diptera: Anthomyiidae). Environ. Entomol. 26: 84-90.

Johnsen, S; Gutierrez, A P; Jorgensen, J. 1997. Overwintering in the **cabbage root fly** *Delia radicum*: A dynamic model of temperature-dependent dormancy and post-dormancy development. J. Appl. Ecol. 34: 21-28.

**Cassava**

Gutierrez, A. P., B. Wermelinger, F. Schulthess, J. U. Baumgartner, J. S. Yaninek, H. R. Herren, P. Neuenschwander, B. Lohr, W. N. O. Hammond and C. K. Ellis. 1987 An overview of a system model of cassava and cassava pests in Africa. Insect Sci. Applic. 8: 919-924.

Gutierrez, A. P., B. Wermelinger, F. Schulthess, J. U. Baumgärtner, H. R. Herren, C. K. Ellis, and J. S. Yaninek. 1988. Analysis of biological control of cassava pests in Africa: I. Simulation of carbon nitrogen and water dynamics in cassava. J. Appl. Ecol. 25: 901-920

Gutierrez, A. P., J. S. Yaninek, B. Wermelinger, H. R. Herren and C. K. Ellis. 1988 Analysis of biological control of cassava pests in Africa: III. **Cassava green mite** *Mononychellus tanajoa*. J. Appl. Ecol. 25: 941-950.

Gutierrez, A. P., J. S. Yaninek, P. Neuenschwande and C. K. Ellis. 1999 A physiologically based metapopulation dynamics: the tritrophic cassava system as a case study. Ecological Modelling. 123:225-242.

Gutierrez, A. P., J. S. Yaninek, P. Neuenschwande and C. K. Ellis. 2000 A physiologically based tritrophic metapopulation model of the African cassava foodweb. Frustula Entomologica. 123:2138-158.

Gutierrez, A. P., P. Neuenschwander and J.J.M. van Alphen.1993. Factors affecting the establishment of natural enemies: biological control of the **cassava mealybug** in West Africa by introduced parasitoids. J. Appl. Ecol. 30: 706-721.

Gutierrez, A. P., P. Neuenschwander, F. Schulthess, H. R. Herren, J. U. Baumgärtner, B. Wermelinger, J. S. Yaninek and C. K. Ellis. 1988 Analysis of biological control of cassava pests in Africa: II. **Cassava mealybug** *Phenacoccus manihoti*. J. Appl. Ecol. 25: 921-940

Yaninek, J. S., H. R. Herren and A. P. Gutierrez. 1987 The biological basis for the seasonal outbreak of **cassava green mites** in Africa. Insect Sci. Applic. 8: 861-865.

**Coffee**

Cure, J.R., D. Rodríguez, A.P. Gutierrez, and L. Ponti (2020) The coffee agroecosystem: bio-economic analysis of coffee berry borer control (*Hypothenemus hampei*). Nature - Scientific Reports, 10: 12262. https://doi.org/10.1038/s41598-020-68989-x [Open Access]

Cure, J.R., H.S. Dos Santos, J.C. Moraes, E. F. Vilela and A. P. Gutierrez. 1988. Fenologia e Dinamica Populacional da Broca do café *Hypothenemus hampei* (Ferrari)relacionadas as fases de desenvolvimento do fruto. . Ann. Entomol. Soc. Brazil. 27 (3): 325 —335.

Cure, José Ricardo, Daniel Rodríguez, Andrew Paul Gutierrez, Luigi Ponti **(2019**) A coffee agroecosystem model: Bioeconomics of coffee berry borer control (*Hypothenemus hampei*). Nature - Scientific Reports | (2020) 10:12262 | https://doi.org/10.1038/s41598-020-68989-x

Gutierrez, A. P., A. Villacorta, J.R. Cure and C. K. Ellis. 1998. A tritophic analysis of the coffee (*Coffee arabica*) - coffee berryborer (*Hypothenemus hampei* (Ferrari)) - parasitoid system. Ann. Entomol. Soc. Brazil. 27 (3): 358 —385.

Rodríguez, D., Cure J.R., Gutierrez A.P., and Cotes J.M. (2017) A coffee agroecosystem model: III. Parasitoids of the coffee berry borer (*Hypothenemus hampei*). *Ecological Modelling* 363: 96-110. https://doi.org/10.1016/j.ecolmodel.2017.08.008

Rodríguez, D., J.R. Cure, J.M. Cotes, A.P. Gutierrez, and F. Cantor (2013) A coffee agroecosystem model: II. Dynamics of coffee berry borer, *Ecological Modelling* 248:203-214. http://doi.org/10.1016/j.ecolmodel.2012.09.015

Rodríguez, Daniel, José Ricardo Cure, José Miguel Cotes, Andrew Paul Gutierrez, Fernando Cantor (2011) A coffee agroecosystem model I. Growth and development of the coffee plant. Ecol. modelling 222:3626– 3639.

**Cotton**

Gutierrez A.P., Ponti L., Karanthi K.R., Baumgärtner J., Boggia A., Cure J.R., Rodríguez D., and G. Gilioli (2020) Bio-economics of Indian hybrid Bt cotton and farmer suicides. Environmental Sciences Europe 32:139. https://doi.org/10.1186/s12302-020-00406-6 [Open Access]

Gutierrez, A. P and S. Ponsard, 2006. Physiologically based model of *Bt* cotton-pest interactions: I. **Pink bollworm**: resistance, refuges, and risk. Ecological Modelling 191:346-359.

Gutierrez, A. P., G. Butler, Y. Wang, and D. Westphal. 1977. The interaction of the **pink bollworm**, cotton, and weather. Can. Entomol. 109: 1457-1468.

Gutierrez, A. P., J.J. Adamcyzk Jr. and S. Ponsard. 2006. A Physiologically based model of *Bt* cotton-pest interactions: II. **bollworm-defoliator-natural enemy** interactions. Ecological Modelling 191: 360-382.

Gutierrez, A. P., L. A. Falcon, W. B. Loew, P. Leipzig, and R. van den Bosch. 1975. An analysis of cotton production in California: A model for Acala cotton and the efficiency of defoliators on its yields. Env. Ent. 4(1): 125-136.

Gutierrez, A. P., M. A. Pizzamiglio, W. J. Dos Santos, R. Tennyson and A. M.Villacorta. 1984. A general distributed delay time varying life table plant population model: cotton (*Gossypium hirsutum* L.) growth and development as an example. Ecol. Modelling 26: 231-249.

Gutierrez, A. P., T. F. Leigh, Y. Wang, and R. D. Cave. 1977. An analysis of cotton production in California: ***Lygus hesperus*** injury--an evaluation. Can. Entomol. 109: 1375-1386.

Gutierrez, A. P., W. J. Dos Santos, A. Villacorta, M. A. Pizzamiglio, C. K. Ellis, L. H. Carvalho and N. D. Stone. 1991. Modelling the interaction of cotton and the cotton **bolll weevil**. I. A comparison of growth and development of cotton varieties. J. Appl. Ecol. 28: 371-397.

Gutierrez, A. P., W. J. Dos Santos, M. A. Pizzamiglio, A. M. Villacorta, C. K. Ellis, C.A.P. Fernandes and I. Tutida. 1991. Modelling the interaction of cotton and the cotton **boll weevil**. II. Boll weevil (*Anthonomus grandis*) in Brazil. J. Appl. Ecol. 28: 398-418.

Gutierrez, A. P., Y. H. Wang, and R. Daxl. 1979. The interaction of cotton and **boll weevil** (Coleoptera: Curculionidae) - a study of co-adaptation. Can. Entomol. 111: 357-366.

Gutierrez, A.P. (2018) Hybrid *Bt* cotton: a stranglehold on subsistence farmers in India. Current Science 115(12) doi: 10.18520/cs/v115/i12/2206-2210

Gutierrez, A.P., 2005. Tritrophic effects in Bt cotton. Bull. Sci. Tech. & Society. 25: 3540-360.

Gutierrez, A.P., C.K. Ellis, T. d'Oultremont and Luigi Ponti. 2006. Climatic limits of pink bollworm in Arizona and California: effects of climate warming. Acta Oecologica 30: 353-364.

Gutierrez, A.P., L. Ponti, and J. Baumgartner. 2017. A critique on the paper 'Agricultural biotechnology and crop productivity: macro-level evidence on contribution of Bt cotton in India.' *Current Science* (Proceedings of the National Academy of India) 112: 690-693. http://www.currentscience.ac.in/Volumes/112/04/0690.pdf

Gutierrez, A.P., L. Ponti, H.R. Herren, J.U. Baumgärtner, and P.E. Kenmore (2015) Deconstructing Indian cotton: weather, yields, and suicides. *Environmental Sciences Europe* 27:12 (17pages). http://doi.org/10.1186/s12302-015-0043-8 [Open Access]

Gutierrez, Andrew Paul, Peter E. Kenmore, and Aruna Rodrigues (2019) When biotechnologists lack objectivity. Current Science (National Academy of Science India) 117 (9): 1-8

Gutierrez, Andrew Paul, Peter E. Kenmore, and Luigi Ponti.  Hybrid Bt cotton is failing in India: cautions for Africa. *Environmental Sciences Europe (2023) 35:93* https://doi.org/10.1186/s12302-023-00804-6

Mills, N. J. and A P. Gutierrez, 1999. Prospective modelling in biological control: An analysis of the dynamics of heteronomous hyperparasitism in a cotton-whitefly-parasitoid system. J. Appl. Ecol., 33: 1379-1394.

Stone, N. D., and A. P. Gutierrez. I986 I. A field-oriented simulation of pink bollworm in southwestern desert cotton. Hilgardia 54: 1-24.

Stone, N. D., and A. P. Gutierrez. 1986 II. A management model for pink bollworm in southwestern cotton. Hilgardia 54: 25-41.

Stone, N. D., A. P. Gutierrez, W. M. Getz, and R. Norgaard. 1986 III. Strategies for pink bollworm control in southwestern desert cotton: An economic simulation study. Hilgardia 54:42-56.

Wang, Y., A. P. Gutierrez, G. Oster, and R. Daxl. 1977. General population model for plant growth and development: coupling cotton-herbivore interactions. Can. Entomol. 109: 1359-1374.

**Cowpea**

Tamo, M.J. Baumgärtner and A.P. Gutierrez. II 1993. Modelling the interaction between cowpea and the bean flower thrips *Megalurothrips sjostedti* (Trybom) (Thysanoptera, Thripidae). Ecol. Modelling 70: 89-113.

**Bioeconomic applications**

Baumgärtner, J., Gianni Gilioli, Getachew Tikubet, Andrew Paul Gutierrez. (2008) Eco-social analysis of an East African agropastoral system: management of tsetse and bovine trypanosomiasis. Ecological Economics 65:124-135.

Gutierrez, A. P., U. Regev and H. Shalit. 1979. An economic optimization model of pesticide resistance: Alfalfa and Egyptian **alfalfa weevil** - an example. Environ. Entomol. 8: 101-107.

Gutierrez, A. P., Y. Wang, and U. Regev. 1979. An optimization model for *Lygus hesperus* (Heteroptera: Miridae) damage in cotton: The economic threshold revisited. Can. Entomol. 111: 41-54.

Gutierrez, A.P., Regev, U. 2005. The bioeconomics of tritrophic systems: applications to invasive species. Ecological Economics 52:382-396.

Neuenschwander, P., W.N.O. Hammond, A. P. Gutierrez, A. R. Cudjoe, R. Adjakloe, J. U. Baumgärtner and U. Regev. 1989. Impact assessment of the biological control of the cassava mealybug, *Phenacoccus manihoti* Matile-Ferrero (Hemiptera: Pseudococcidae), by the introduced parasitoid *Epidinocarsis lopezi* (De Santis) (Hymenoptera: Encyrtidae). Bull. Ent. Res. 79: 579-594.

Pemsl, D., Gutierrez, A.P., Waibel, H. (2007). The Economics of Biotechnology under Ecosystems Disruption. Ecological Economics. 66:177-183.

Pemsl, D., H. Waibel and A.P. Gutierrez. 2005. Why do some Bt-cotton farmers in China continue to use high levels of pesticides? International J. Agric. Sustainability 3:44-56.

Pemsl, D., Waibel H., and A. Gutierrez (2003): Productivity analysis of Bt cotton: A case study from Shandong province, China. Paper presented at the 7th ICABR Conference on Public Goods and Public Policy for Agricultural Biotechnology. Ravello, Italy, June 29 to July 3.

Regev, U., A. P. Gutierrez, J. E. DeVay and C. K. Ellis. 1990 Optimal strategies for management of **verticillium wilt**. Agric. Syst. 33: 139-152.

Regev, U., A. P. Gutierrez, S. J. Schreiber, and D. Zilberman. 1998. Biological and Economic Foundations of Renewable Resource Exploitation. Ecological Economics, 26: 227-242.

**Fruit flies**

Gutierrez, A.P. and Luigi Ponti (2011). Assessing the invasive potential of the **Mediterranean fruit fly** in California and Italy. Biol. Invasion DOI 10.1007/s10530-011-9937.

Gutierrez, A.P., L. Ponti, and G. Gilioli (2014) Comments on the concept of ultra-low, cryptic tropical fruit fly populations. *Proceedings of the Royal Society B: Biological Sciences* 281: 20132825. http://doi.org/10.1098/rspb.2013.2825 [Freely Available]

Gutierrez, A.P., L. Ponti, M., J. Suckling, M. Neteler (2021) Invasive potential of **tropical fruit flies** in temperate regions under climate change. Nature - Biological communications 4:1141 | https://doi.org/10.1038/s42003-021-02599-9.

**Grape**

Baumgärtner, J., A.P. Gutierrez, S. Pesolillo, and M. Severini (2012) A model for the overwintering process of **European grapevine moth** *Lobesia botrana* (Denis & Schiffermüller) (Lepidoptera, Tortricidae) populations. *Journal of Entomological and Acarological Research* 44:e2. http://doi.org/10.4081/jear.2012.e2

Gutierrez A.P., Ponti L., Cooper M.L., Gilioli G., Baumgärtner J., Duso C., 2012. Prospective analysis of the invasive potential of the **European grapevine moth** *Lobesia botrana* (Den. & Schiff.) in California. *Agricultural and Forest Entomology*, 14: 225-238. http://dx.doi.org/10.1111/j.1461-9563.2011.00566.x

Gutierrez, A. P., D. W. Williams, and H. Kido. 1985. A model of **grape** growth and development: the mathematical structure and biological considerations. Crop Sci. 25: 721-728.

Gutierrez, A.P., K.M. Daane, Luigi Ponti, Vaughn M. Walton, C.K. Ellis, (2007) Prospective evaluation of the biological control of the **vine mealybug**: refuge effects. J. Appl. Ecol. 44: 1-13.

Gutierrez, A.P., L. Ponti, J. Baumgärtner, and G. Gilioli (2017) Climate warming effects on grape and grapevine moth (*Lobesia botrana*) in the Palearctic region. *Agricultural and Forest Entomology*, http://doi.org/10.1111/afe.12256

Gutierrez, Andrew Paul, Luigi Ponti, Mark Hoddle, Rodrigo P.P. Almeida, and Nicola A. Irvin (2011). Evaluation of the factors affecting the geographic distribution and abundance of **glassy-winged sharpshooter** (*Homalodisca vitripennis* Germar) and two egg parasitoids in California-- Environ. Entomol. 40(4):755-769.

Ponti, L., A.P. Gutierrez, A. Boggia, and M. Neteler (2018) Analysis of grape production in the face of climate change. Climate 6:20. https://doi.org/10.3390/cli6020020 [Open Access]

Wermelinger, B. , J. Baumgärtner and A.P. Gutierrez. 1991. A demographic model of assimilation and allocation of carbon and nitrogen in grapevines. Ecol. Modelling 53: 1-26.

**Light brown apple moth**

Gutierrez, A.P., N. J. Mills and L. Ponti (2010.) Limits to the potential distribution of **light brown apple** moth in Arizona–California based on climate suitability and host plant availability. Bio. Invasions. DOI 10.1007/s10530-010-9725-8

**Mosquitoes**

Gilioli, G., Pasquali, S., Ponti, L., Calvitti, M., Moretti, R., & Gutierrez, A. P. (2015) Modelling the potential distribution and abundance of ***Aedes albopictus*** in Europe in light of the climate change scenario. *Conference: Impacts of Environmental Changes on Infectious Diseases*, 23-25 March 2015, Sitges, Spain.

Pasquali, S., L. Mariani, M. Calvitti, R. Moretti, L. Ponti, M. Chiari, G. Sperandio, and G. Gilioli. 2020. Development and calibration of a model for the potential establishment and impact of ***Aedes albopictus*** in Europe. Acta Tropica 202:doi:10.1016/j.actatropica.2019.105228.

**Oleander scale**

Gutierrez, A.P., M.A. Pizzamiglio (2007) A Regional Analysis of Weather Mediated Competition between a Parasitoid and a Coccinellid Predator of **Oleander Scale**. *Neotropical Entomology* 36(1):70-83.

**Olive**

Di Paola, Arianna, Edmondo Di Giuseppe, Andrew Paul Gutierrez, Luigi Ponti, Massimiliano Pasqui (2023) Climate stressors modulate interannual olive yield at province level in Italy: a composite index approach to support crop management . J. Agronomy and Crop Science, 00, 1-14 https://doi.org/10.1111/jac.12636

Gutierrez, Andrew Paul, Luigi Ponti and Q. A. Cossu (2009) Prospective comparative analysis of global warming effects on olive and **olive fly** (*Bactrocera oleae* (Gmelin)) in Arizona-California and Italy. Climatic Change 95:195-217.

Ponti, L., A.P. Gutierrez, P.M. Ruti, and A. Dell'Aquila (2014) Fine-scale ecological and economic assessment of climate change on olive in the Mediterranean Basin reveals winners and losers. *PNAS Proceedings of the National Academy of Sciences* 111:5598-5603. http://doi.org/10.1073/pnas.1314437111 [Open Access]

Ponti, L., Q.A. Cossu, and A.P. Gutierrez (2009) Climate warming effects on the olive-*Bactrocera oleae* system on Mediterranean Islands: Sardinia as an example. Global Climate Change. 15(12)2:874-2884 http://dx.doi.org/10.1111/j.1365-2486.2009.01938.x

Rochat, J. and A. P. Gutierrez (2001) Weather mediated regulation of **olive scale** by two parasitoids. J. Anim. Ecol. 70: 476-490.

**Rice**

d' Oultremont, T. and A.P. Gutierrez. 2002. A multitrophic model of a **rice-fish** agroecosystem: II. Linking the flooded rice-fishpond systems, Ecological Modelling 155 (2-3):159 - 176.

d' Oultremont, T. and A.P. Gutierrez. 2002. A multitrophic model of a **rice-fish** agroecosystem: I. A tropical fishpond food web, Ecological Modelling 156 (2-3): 123 - 142.

Graf, B., A. P. Gutierrez, O. Rakotobe, P. Zahner and V. Delucchi. 1990 A simulation model for the dynamics of rice growth and development. II. Competition with weeds for nitrogen and light. Agricultural Systems 32: 367-392.

Graf, B., J. Baumgärtner and A. P. Gutierrez. 1990. Modeling agroecosystem dynamics with the metabolic pool approach. Bulletin de la Societe Entomologique Suisse 63: 465-476.

Graf, B., O. Rakotobe, P. Zahner, V. Delucchi and A. P. Gutierrez. 1990 A simulation model for the dynamics of rice growth and development. I. The carbon balance. Agricultural Systems 32: 341-365.

**Screwworm**

Gutierrez, A.P., L. Ponti, and P.A. Arias. 2019. Deconstructing the eradication of **new world screwworm** in North America: retrospective analysis and climate warming effects. Medical and Veterinary Entomology, https://doi.org/10.1111/mve.12362 [Open Access]

Gutierrez, A.P., and L. Ponti (2014) The **new world screwworm**: prospective distribution and role of weather in eradication. *Agricultural and Forest Entomology* 16:158-173. http://doi.org/10.1111/afe.12046

**Spotted winged drosophila.**

Asplen, M.K., G. Anfora, A. Biondi, D.-S. Choi, D. Chu, K.M. Daane, P. Gibert, A. P. Gutierrez, K.A. Hoelmer, W.D. Hutchison, R. Isaacs, Z.-L. Jiang, Z. Kárpáti, M. T. Kimura, M. Pascual, C. R. Philips, C. Plantamp, L. Ponti, G. Vétek, H. Vogt, V.M. Walton, Y. Yu, L. Zappalà, and N. Desneux (2015) Invasion biology of **spotted wing Drosophila** (Drosophila suzukii): a global perspective and future priorities. Journal of Pest Science 88:469-494. http://doi.org/10.1007/s10340-015-0681-z [PDF Open Access]

Gutierrez, A.P., L. Ponti, and D.T. Dalton. 2016. Analysis of the invasiveness of **spotted wing drosophila** (Drosophila suzukii) in North America, Europe, and the Mediterranean Basin. Biological Invasions 18:3647-3663. http://doi.org/10.1007/s10530-016-1255-6 [PDF Free to View]

**Theory**

Baumgärtner, J. U., A. P. Gutierrez, and A. Klay. 1988 Elements of modeling the dynamics of tritrophic population interactions. Exp. Appl. Acarology 5: 243-263.

Graf, B., J. Baumgärtner and A. P. Gutierrez. 1990. Modeling agroecosystem dynamics with the metabolic pool approach. Bulletin de la Societe Entomologique Suisse 63: 465-476.

Gutierrez, A.P. 1992. The physiological basis of ratio dependent theory. Ecology 73:1529-53.

Gutierrez, A.P., and L. Ponti. 2013. Eradication of invasive species: why the biology matters. *Environmental Entomology* 42(3): 395-411. http://dx.doi.org/10.1603/EN12018 [Open Access]

Gutierrez, A.P., N.J. Mills, S.J. Schreiber, and C.K. Ellis. 1994. A Physiologically Based Tritrophic Perspective on Bottom-Up - Top-Down Regulation of Populations. Ecology 75: 2227-2242.

Schreiber ,S.J., and A.P.Gutierrez (1999). Insect Invasions and Community Assembly**.** In *Ecological Entomology*, Huffaker, C.B. and A. P. Gutierrez (editors). John Wiley and Sons.

Schreiber, S.J., and A. P. Gutierrez. 1998. A supply-demand perspective of species invasions and coexistence: applications to biological control. Ecol. Modelling 106: 27-45.

Schreiber, S.J., N.J. Mills and A.P. Gutierrez (2001) Host limited Dynamics of Autoparasitoids. J. Theor. Biol. 212 (2): 141-153.

Wang, Y., and A. P. Gutierrez. 1980. An assessment of the use of stability analysis in population ecology. J. Anim. Ecol. 49: 435-452.

**Thimble berry aphid**

Gilbert, N. E., and A. P. Gutierrez. 1973. A plant-aphid-parasite relationship. J. Anim. Ecol. (42): 323-340.

**Tomato**

Wilson, L. T., A. P. Gutierrez, R. Tennyson, and F. G. Zalom. 1986. A physiological based model for processing tomatoes: crop and pest management. 2nd Intern. Symp. on Proc. Tomatoes, U.C. Davis (Aug. 18-19, 1986). Acta Hort. 200: 125-132.

**Tomato pinworm**

Ponti, L., and Gutierrez, An.P. 2023 Challenging the status quo in invasive species assessment using mechanistic physiologically based demographic modeling. Environment, Development and Sustainability https://doi.org/10.1007/s10668-023-03698-9

Ponti, L., A.P. Gutierrez, M.R. de Campos, N. Desneux, A. Biondi, M. Neteler. 2021. Biological invasion risk assessment of *Tuta absoluta*: mechanistic versus correlative methods. Biol Invasions. https://doi.org/10.1007/s10530-021-02613-5

**Verticillium wilt in cotton**

Devay, J.E., A.P. Gutierrez, G.S. Pullman, R.J.Wakeman, R.H. Garber, D.P. Jeffers, S.N. Smith, P.B. Goodell, and P.A. Roberts. 1997. Inoculum densities of *Fusarium oxysporum f. sp. vasinfectum* and *Meloidogyne incognita* in relation to the development of fusarium wilt and the phenology of cotton plants (*Gossypium hirsutum*). Phytopath. 87: 341-346.

Gutierrez, A. P., J. E. Devay, G. S. Pullman, and G. E. Friebertshauser. 1982. A model of verticillium wilt in relation to cotton growth and development. Phytopath. 73: 89-95. 1983

**Veterinary**

Gilioli G., M. Groppi, M.P. Vesperoni, J. Baumgärtner, and A.P.  Gutierrez. 2009. An epidemiological model of **East Coast Fever** in African livestock. Ecol. Modelling 220:1652-1662.

Gutierrez, A.P., G. Gilioli and J. Baumgärtner. 2009 Eco-social Consequences of Disease Management in East African Agro-Pastoral Systems: Policy Implications. Proceedings of the National Academy of Science 106: 13136-13141.

**Yellow starthistle**

Gutierrez, A.P., L. Ponti, M. Cristofaro, L. Smith, and M.J. Pitcairn. 2017. Assessing the biological control of **yellow starthistle** (*Centaurea solstitialis* L): prospective analysis of the impact of the rosette weevil (*Ceratapion basicorne* (Illiger)). Agricultural and Forest Entomology 19: 257- 273 http://doi.org/10.1111/afe.12205 [PDF Free to View]

Gutierrez, A.P., M.J. Pitcairn, C.K. Ellis, N. Carruthers, and R. Ghezelbash. 2005. Evaluating biological control of **yellow starthistle** (*Centaurea solstitialis*) in California: A GIS based supply–demand demographic model. Biological Control 34: 115-131.

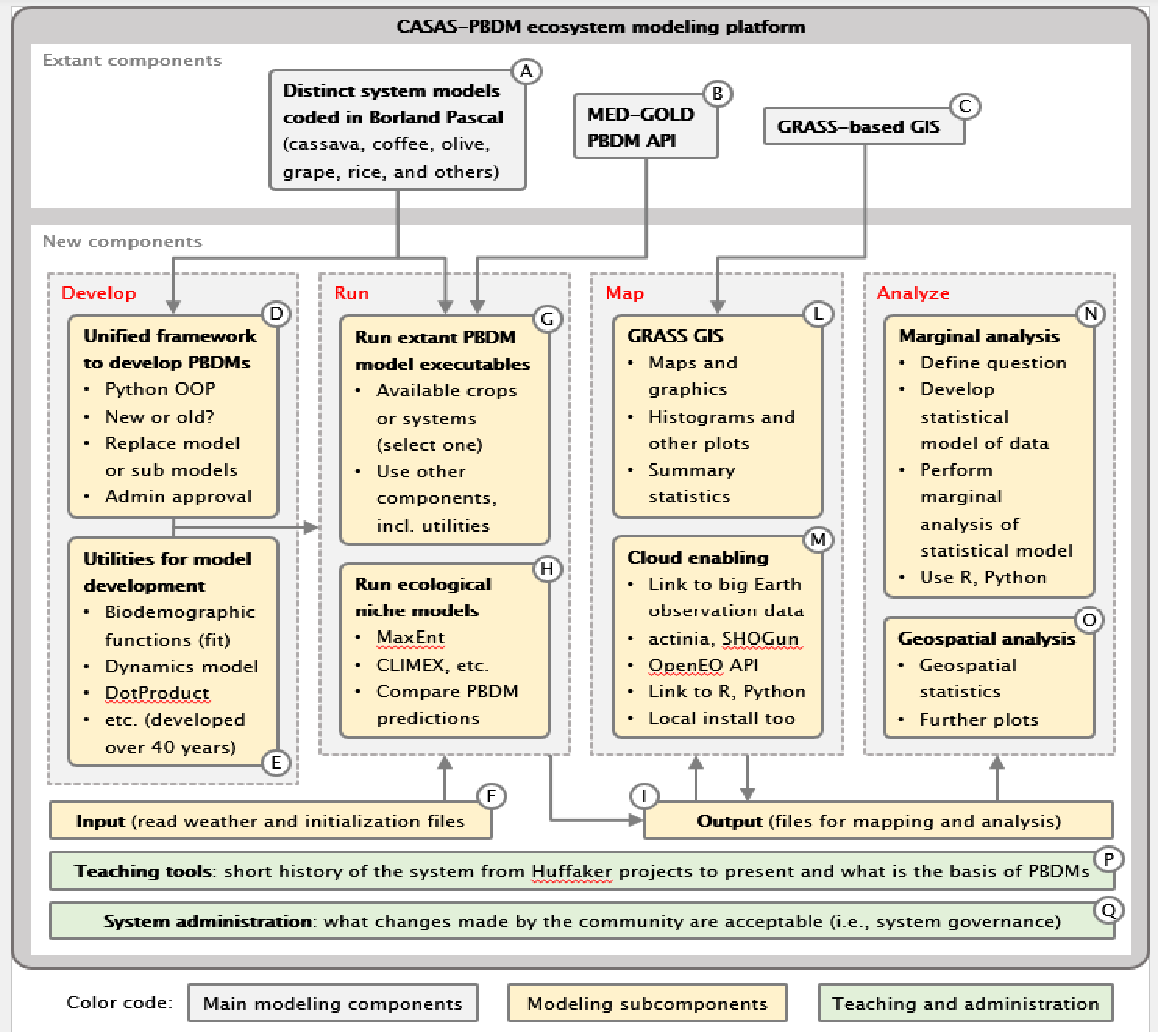


Supplemental Materials **Figure 1.** Platform for implementing the CASASGlobal physiologically based demographic models (PBDMs) platform. **(A)** Distinct system models coded in Borland Pascal; **(B)** MED-GOLD PBDM application programming interface (API); **(C)** GRASS-based GIS; **(D)** Generalized framework for PBDM development; **(E)** Utilities for model development; **(F)** Input (read weather and setup files); **(G)** Run extant PBDM model executables; **(H)** Run ecological niche models; **(I)** Output (files for mapping and analysis); **(L)** GRASS GIS; **(M)** Cloud enabling (e.g., actinia, SHOGun, OpenEO); **(N)** Marginal analysis; **(O)** Geospatial analysis; **(P)** Teaching tools: short history of the system and what is the basis of PBDMs; **(Q)** System administration: what changes are acceptable (i.e., open-source system governance.)

https://ec.europa.eu/info/funding-tenders/opportunities/portal/screen/opportunities/horizon-results-platform/32534
